# Cell type-specific contributions to impaired blood-brain barrier and cerebral metabolism in presymptomatic 5XFAD mice

**DOI:** 10.1101/2025.04.23.650260

**Authors:** Minmin Yao, Na Sun, Raleigh Linville, Zhiliang Wei, Aaron Kakazu, Yuxiao Ouyang, Ruoxuan Li, Lida Du, Haitong Wang, Yuan Zhou, Yanli Jiang, Ziqin Zhang, Anna Li, Hanzhang Lu, Jiadi Xu, Manolis Kellis, Myriam Heiman, Wenzhen Duan

## Abstract

Altered cerebral metabolism and blood-brain barrier (BBB) dysfunction are emerging as critical contributors to the preclinical phase of Alzheimer’s disease (AD), underscoring their role in early pathogenesis. To identify sensitive biomarkers before irreversible neuronal loss and cognitive decline, we examined 5XFAD mice at 3 months of age by applying multiple advanced MRI techniques. Arterial spin tagging based MRI revealed increased BBB permeability and water extraction fraction, indicating compromised BBB integrity at the early stage of pathogenesis in 5×FAD mice. Despite preserved cerebral blood flow, a decreased unit mass cerebral metabolic rate of oxygen (CMRO2) was evident in the same cohorts of 5XFAD mice. Interestingly, a region-specific decrease of tissue pH values was detected in the hippocampus of these 5XFAD mice by creatine chemical exchange saturation transfer MRI. Elevated neuronal H4K12 lactylation in the hippocampus supports the reduced pH values. To further dissect the cellular and molecular mechanisms underlying these MRI-detectable changes in 5XFAD mice, we conducted single-nucleus RNA sequencing (snRNA-Seq) with optimized blood vessel enrichment protocols. Our results revealed cell type-specific transcriptomic changes in the hippocampus of 3-month-old 5XFAD mice, including downregulation of synaptogenesis and synaptic transmission genes in the CA1 and dentate gyrus excitatory neurons, impaired endothelial gene expression linked to brain barrier function and angiogenesis, altered innate immune response genes in astrocytes, as well as upregulation of cholesterol biosynthesis and metabolism genes in the CA1 excitatory neurons. These findings underlie the intricate interplay between BBB disruption and metabolic dysregulation before the onset of cognitive decline in AD. Our study demonstrates that BBB dysfunction and cerebral metabolic alterations preceded brain hypoperfusion and cognitive decline, emphasizing potential molecular pathways for early intervention. These findings, once validated in human studies, could significantly enhance early diagnosis and inform novel therapeutic strategies targeting early AD pathogenesis.

## Introduction

Alzheimer’s disease (AD) is characterized by a prolonged preclinical phase, spanning approximately two decades, during which pathology silently accumulates without overt clinical symptoms^1^. Despite tremendous advancements in understanding AD pathogenesis, effective treatments that slow or prevent clinical progression remain elusive. Disease pathology begins decades before symptom onset, making early intervention crucial but challenging. By the time cognitive symptoms emerge, the disease is already in an advanced state. Moreover, while approaches to remove amyloid β (Aβ) plaques have been developed, they have shown limited efficacy in addressing other major drives of cognitive decline, including tau pathology and neuroimmune dysfunction. Thus, identifying reversible functional changes early in the disease process is critical to developing disease-modifying therapies and curbing the projected rise in global AD prevalence to 150 million by 2050.

A growing body of experimental clinical evidence shows that age-related blood-brain barrier (BBB) disruption is a key contributor to cognitive decline in AD^2,3^. BBB dysfunction is evident in AD pathogenesis and is suspected to play a causative role^2,4–6^. Animal studies have further elucidated that BBB disruption precedes and exacerbates neuroinflammation, synaptic dysfunction, and cognitive impairment ^7–9^. The leakage of serum factors through a compromised BBB activates microglia, initiating processes such as synaptic over pruning and neuroinflammation that can amplify AD pathology^10–12^. Importantly, preclinical models have shown that strategies to preserve or restore BBB integrity can mitigate cognitive deficits^13,14^, underscoring its potential as a therapeutic target. These lines of evidence indicate the significance of sensitive and non-invasive measurement of BBB permeability in the early stages of AD development.

Cerebral metabolism, another critical aspect of AD pathology, is profoundly altered during the initial stages of AD. The brain has a high energy demand that is met through glucose metabolism, with aerobic glycolysis serving as a complementary pathway to oxidative phosphorylation. Reduction of glucose utilization is a well-documented phenomenon in AD and occurs early in the disease process^15^. Notably, reductions in glucose consumption exceed decreases in blood flow and oxygen utilization, suggesting that disruptions in aerobic glycolysis are among the earliest metabolic changes in AD^16,17^. Evidence from human studies indicates that regions with high resting-state aerobic glycolysis correlate spatially with regions of Aβ deposition, further implicating altered glucose metabolism in early AD pathology. Additionally, under stress conditions, cells may shift from oxidative phosphorylation to aerobic glycolysis, a metabolic reprogramming that supports rapid ATP production and the generation of glycolytic intermediates critical for immune responses^18,19,20^.

In this study, we investigated early-stage changes in the brains of 5XFAD mice^21^, a well-established model of AD, focusing on BBB integrity and cerebral metabolism. We observed compromised BBB integrity and declined cerebral metabolic rate of oxygen (CMRO2) in 3-month-old 5XFAD mice, accompanied by decreased cerebral pH levels in the hippocampus. These changes occurred at an early pathogenic stage prior to cognitive decline and motor behavioral changes^22^, highlighting their potential role as early non-invasive biomarkers in AD progression. Furthermore, we identified an increase in the lactylation of lysine 12 on histone H4 (H4K12La) in hippocampal neurons, a marker associated with metabolic reprogramming and immune function.

To further elucidate the molecular underpinnings of these changes, we performed single-nucleus RNA sequencing (snRNA-seq) of the hippocampus, employing optimized blood vessel enrichment protocols. Our analysis revealed distinct cell type-specific gene expression changes in the hippocampus of 3-month-old 5XFAD mice; endothelial cells exhibited downregulation of BBB-related genes and upregulation of genes associated with immune responses and neuroinflammation, while neurons (particularly excitatory neurons) exhibited upregulation of glycolysis-related and hypometabolism-associated genes. Intriguingly, genes involved in cholesterol biosynthesis and metabolism were upregulated in both excitatory neurons and endothelial cells, further suggesting a potential link between lipid metabolism and early AD pathology.

These findings underscore the importance of BBB integrity and cerebral metabolism as early and potentially reversible contributors to AD pathogenesis. Non-invasive MRI measures of BBB dysfunction and metabolic alterations could serve as valuable diagnostic biomarkers, enabling earlier intervention and facilitating the development of disease-modifying therapies for AD.

## Results

### Compromised BBB integrity precedes extracellular β-amyloid deposits in the hippocampus and cognitive impairment in 5XFAD mice

The BBB controls the molecular exchange between the brain parenchyma and blood, allows the selective removal of metabolic waste from the brain, plays a major role in regulating cerebral blood flow, and connects the central nervous system to blood circulation. BBB disruption is a key factor in the development of neuroinflammation and cognitive impairment ^7–9,23^. BBB breakdown can lead to the permeation of harmful perivascular factors and immune cells into the brain, aggravating neuroinflammation and neuronal death. At the same time, abnormalities in BBB transport systems disrupt the uptake of nutrients such as glucose from the circulation and the clearance of toxic brain-derived proteins like Aβ, further affecting neuronal activity and increasing Aβ burden.

To investigate BBB functional integrity, we employed WEPCAST MRI, a non-invasive, sensitive, and human applicable MR imaging technique that enables the assessment of BBB permeability to water molecules^24,25^. From the magnetically labeled water signal observed at the venous side, the amount of water extravasated across the BBB can be measured. By comparing the WEPCAST MRI results with histological measures of tight junction proteins, we have validated WEPCAST MRI as a sensitive measure of BBB permeability ^24^. Using this technique, we found that patients with mild cognitive impairment manifested an increased BBB permeability to water which correlated with the burden of amyloid and phosphorylated tau ^26^, suggesting that BBB breakdown occurs at the early stage of AD. Interestingly, we detected increased BBB water permeability in 3-month-old 5XFAD mice (Figure 1A), indicated by increased water extraction fraction to water (Mr. 18Da) (Figure 1B) and permeability surface area product (PS) (Figure 1C). When we administered EZ-link^TM^ sulfo-NHS-LC-biotin (biotin; 500 mg/kg, MW 556.59 Da) to the 5XFAD mice, we did not detect biotin linkage outside blood vessels (Figure 1D), suggesting that the early BBB permeability increase in 5XFAD mice is subtle and only to small molecules like water, and demonstrated the sensitivity of WEPCAST MRI in measuring BBB permeability changes. These results may explain prior findings that showed preserved BBB permeability in 5XFAD mice when using more traditional approaches^27^. Upon histological examination of the 5XFAD mouse brain, subtle Thioflavin S positive Aβ fibrils were evident in the frontal cortex and subiculum area **(Figure 1E),** while other brain regions were free from visible Aβ deposition at 3 months of age. Notably, these mice are presymptomatic as they show normal cognitive function in both the Y-maze (**Figure 1F**) and novel object recognition tests **(Figure 1G-H).**

**Figure 1.**
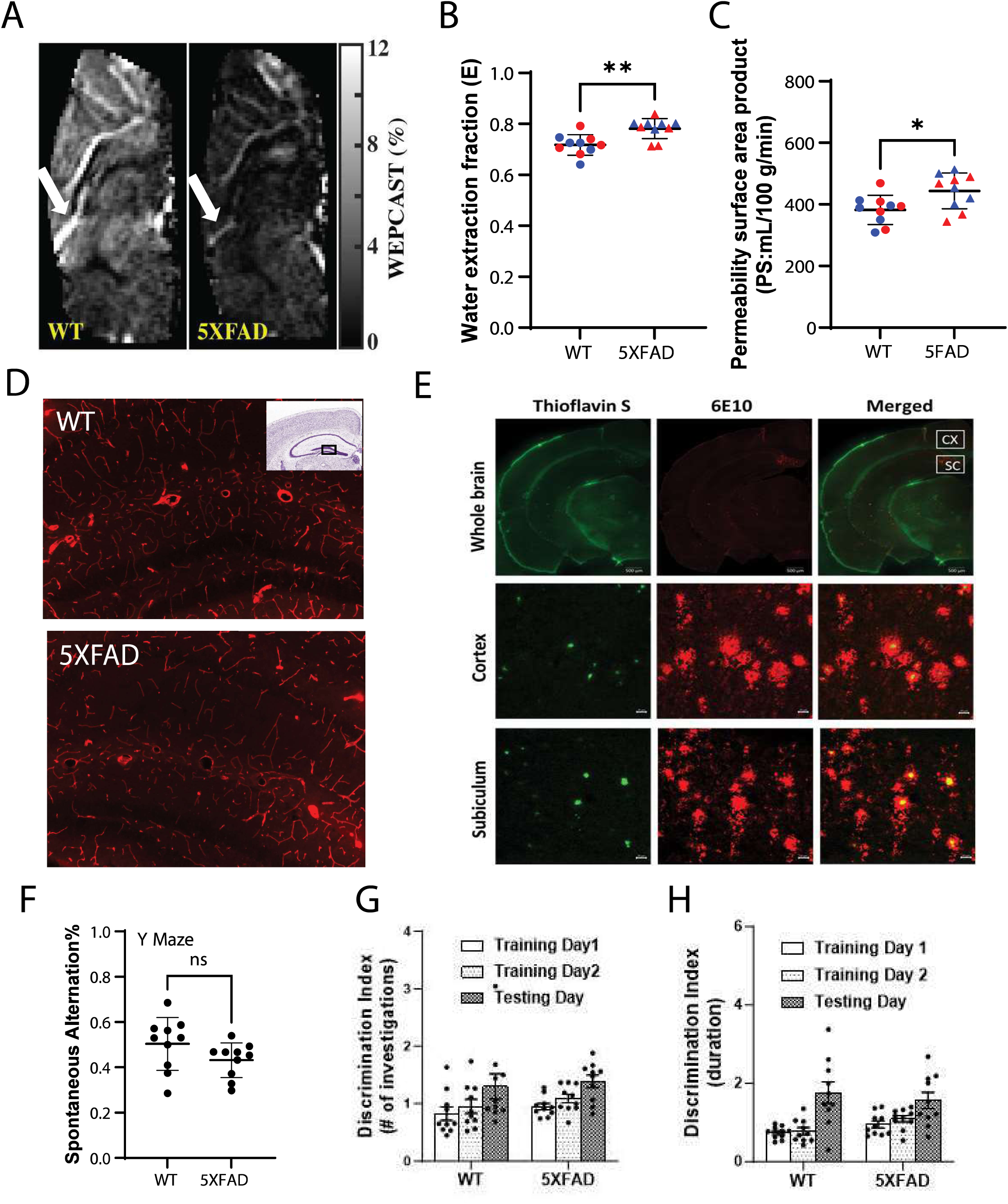
Increased blood-brain barrier permeability precedes amyloid beta fibrillation and cognitive behavioral deficits in presymptomatic 5XFAD mice. (A) Representative WEPCAST MRI images from 5XFAD and age-matched wild-type (WT) mouse brains highlight differences in water extraction at the Great Vein of Galen (GVG). Brighter regions indicate higher levels of magnetically labeled water, which are reduced in 5XFAD brains. (B) Quantified water extraction fraction shows increased BBB permeability in 5XFAD mice. n= 10 mice/group. Male and female mice were equally included. **p<0.01 between WT and 5XFAD groups by two-tailed Student’s t-test. Red and blue color points represent female and male data, respectively. (C) BBB permeability is demonstrated by the permeability surface area product (PS) which is derived from cerebral blood flow and the water extraction fraction. **p<0.01 between WT and 5XFAD groups by two-tailed Student’s t-test. (D) Tight junction integrity was assessed using EZ-link™ Sulfo-NHS-LC-Biotin as a barrier tracer. (E) Amyloid β deposition was visualized in 3-month-old 5XFAD mice via Thioflavin S (Aβ fibrils) and 6E10 antibody staining in the cortex (CX) and subiculum (SC). Scale bars: 500 µm (upper panels); 10 µm (lower panels). (F-H) Behavioral analysis in 3-month-old mice, including Y-maze and novel object recognition tests, revealed no significant differences in spatial memory or cognitive function between 5XFAD and WT mice (n=10 per group, 5 male and 5 female mice/group).

Furthermore, we evaluated the inflammation and morphology of cerebral blood vessels in 3-month-old 5XFAD mice. We observed increased levels of glial fibrillary acidic protein (GFAP) **(Figure S1A-B),** indicating astrocyte activation. While there were no significant changes in ionized calcium-binding adaptor molecule 1 (Iba1) protein levels in Western blotting analysis **(Figure S1C upper** panel, D), microglia in the 5XFAD mice exhibited a reactive morphology **(Figure S1C lower** panel), indicating an early stage of microglia activation. The expression of the tight junction protein claudin-5 showed no significant differences in protein levels **(Figure S1E-F).** In addition, the intensities of the pericyte marker CD13 **(Figure S2)** and the endothelial cell marker CD31 **(Figure S3)** were unchanged, indicating that pericyte and endothelial cell populations were unaffected morphologically at this stage of 5XFAD mice. Additionally, aquaporin-4 (Aqp4) polarization **(Figure S3)** in both the cortex and hippocampus remained unchanged, suggesting that water channel distribution was maintained.

In summary, our comprehensive assessment of cellular and molecular components of the neurovascular unit in 3-month-old 5XFAD mice reveals significant astrocyte activation and mild reactive microglia, while the morphology of endothelial cells and pericytes remains indistinguishable from that in wild-type mice. These findings provide new insights into the early neurovascular changes in a mouse model recapitulating the genetic and neuropathological features of familial AD. The molecular mechanism leading to impaired BBB permeability to water at this early stage of 5XFAD mice warranted further exploration.

### Decreased CMRO2 and hippocampal pH in presymptomatic 5XFAD mice

The brain has a high demand for oxygen compared to other organs. It utilizes approximately 20% of the body’s total oxygen consumption, making tight regulation of oxygen delivery critical for brain function. The quantification of the brain oxygen extraction fraction (OEF), in conjunction with perfusion imaging for cerebral blood flow (CBF), makes the CMRO2 a key measure of brain hemodynamic function. AD patients show regional hypoperfusion and decreased levels of tissue oxygenation ^28^, as well as capillary dysfunction at an early disease stage, which has been reported to be associated with cognitive symptom severity and neurodegeneration ^29^. In recent years, various biological and medical imaging techniques have been developed to assess cerebral oxygenation in animal models and in humans. Measurements of blood oxygenation level dependent (BOLD) signals and CBF in conjunction with hypercapnic or hypertoxic respiratory challenges have been proposed for measuring the relative change and absolute value of CMRO2 ^30^. However, these methods that infer the regional concentration of oxygen by measuring tissue R2, R2’ or R2* relaxation rates or bulk susceptibility, which is sensitive to paramagnetic compounds such as hemoglobin (Hb), are prone to confounders as non-vascular tissue compartments invariably contribute to the signals or are sensitive to changes in vessel caliber and orientation. This limits their applicability for characterizing brain diseases that involve significant remodeling of the vasculature, such as under AD conditions. To assess whether cerebral metabolism is altered in 5XFAD mice at the early disease phase, we utilized T2 relaxation under spin tagging (TRUST) MRI to evaluate global cerebral metabolism ^31^, as indicated by oxygen extraction fraction (OEF) and CMRO2 in the brain. Our results indicate that both CMRO2 and OEF were significantly reduced in 3-month-old 5XFAD mice, with no sex-dependent difference in both measures **(Figure 2A-B),** suggesting early metabolic alterations in both male and female 5XFAD mice. The low oxygen availability to the brain can influence synaptic transmission and lead to neuronal dysfunction ^32^, and an increase in APP processing, thus aggravating Aβ deposition ^33^.

**Figure 2.**
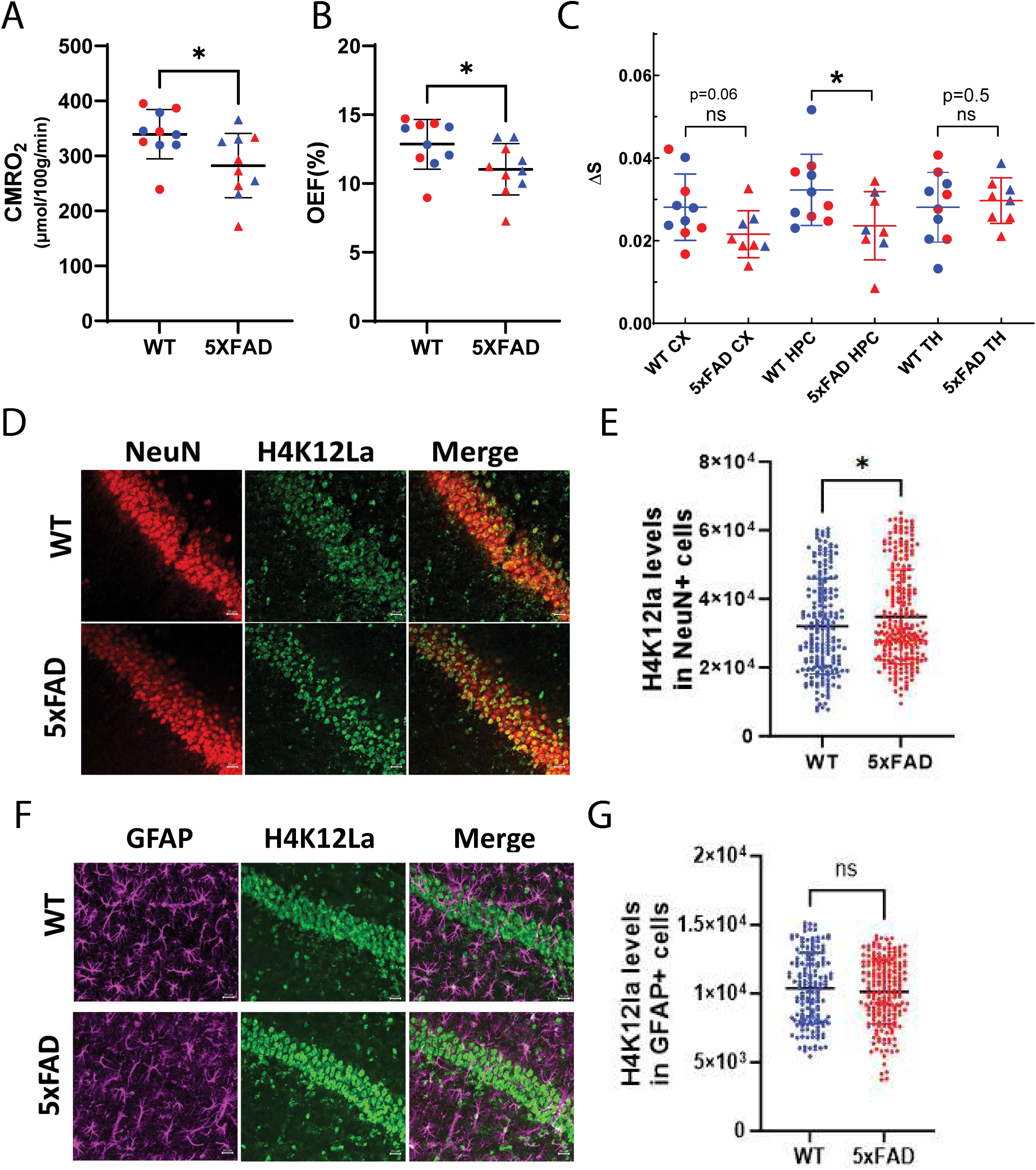
Altered cerebral metabolism in presymptomatic 5XFAD mice. (A) The cerebral metabolic rate of oxygen (CMRO2) was measured in vivo using T2 relaxation under spin tagging (TRUST) and phase-contrast MRI. Data points (n = 10 mice/group, 5 male and 5 female/group) show significant differences (*p < 0.05, two-tailed Student’s t-test). Red dots represent data from female mice, while blue dots represent male mice. (B) Oxygen extraction fraction (OEF) was measured in the brain, revealing significant alterations in 5XFAD mice (n = 10 mice/group, 5 male and 5 female/group), *p < 0.05, two­tailed Student’s t-test). (C) Creatine chemical exchange saturation transfer (CrCEST) MRI detected reduced hippocampal pH levels in 3-month-old 5XFAD mice (n = 10 mice/group, 5 male and 5 female/group), *p < 0.05, two-tailed Student’s t-test). No significant differences were observed in other regions (ns). (D) Representative images showing H4K12 lactylation in neurons. NeuN (red) marks neurons, and green fluorescence indicates H4K12 lactylation. Scale bar = 20 µm. (E) Quantification of neuronal H4K12 lactylation levels across genotypes (n = 200 neurons from 3 mice/group, *p < 0.05, two­tailed Student’s t-test). (F) Representative images showing H4K12 lactylation in astrocytes. GFAP (purple) labels astrocytes and green fluorescence indicates H4K12 lactylation. Scale bar = 20 µm. (G) Quantification of H4K12 lactylation levels in astrocytes across genotypes (n = 200 GFAP-positive cells from 3 mice/group). No significant differences were observed (ns, two-tailed Student’s t-test).

Signs of extracellular acidosis have been observed in the postmortem brain of AD cases ^34^ and was further confirmed by a large cohort study across multi-institutes ^35^. A recent study shows that *in vivo* acidification of mouse brain tissue increases amyloid plaque deposition ^36^. It has been indicated that the reduced intracellular cerebral pH is a consequence of neuroinflammation involved in the development of AD ^37^. Hence, non-invasive *in vivo* assessment of cerebral pH could be a useful biomarker for differentiating between early AD and healthy controls ^38,39^. To investigate whether alterations in cerebral pH are present in the early stage of the 5XFAD mouse model, we conducted creatine chemical exchange saturation transfer (CrCEST) MRI on 3-month-old 5XFAD mice. CrCEST contrast is sensitive to both the concentration of exchangeable protons and pH, as the exchange rate of creatine guanidinium protons is strongly pH-dependent^40^. We analyzed cerebral pH values in three brain regions: the cerebral cortex, hippocampus, and thalamus. A significantly lower cerebral pH was detected in the hippocampus of 5XFAD mice compared to age-matched controls **(Figure 2C).** There was a trend toward reduced pH in the cerebral cortex of AD mice, while no difference was observed in the thalamus **(Figure 2C).** These findings suggest that pH alterations occur in a brain region-specific manner, and the hippocampus is a particularly vulnerable brain structure thus, such a non-invasive measure could be considered as a biomarker for early detection of AD.

### Elevated H4K12la levels in the neurons of presymptomatic 5XFAD mice

To further understand the molecular basis of hippocampal acidification in 5XFAD mice, we hypothesize that a metabolic switch from aerobic metabolism to anaerobic metabolism may underlie the observed reduction in cerebral pH. Glycolysis produces pyruvate, which can be converted into acetyl-CoA in the presence of sufficient oxygen supply through aerobic metabolism or into lactate under low-oxygen conditions. Combined with our finding on reduced cerebral oxygen metabolism (reduced CMRO2) in the 5XFAD brain, the decreased hippocampal pH value may imply a metabolic switch to increased anaerobic metabolism in the AD mouse brain as a compensatory energy supply under lower oxygen conditions. Anaerobic metabolism produces lactate and increased lactate levels can promote histone lactylation, including H4K12 lactylation, which can regulate gene expression in response to metabolic changes ^41^. Previous studies have reported increased H4K12 lactylation in 12-month-old 5XFAD mice^42^ and late-stage AD patients. Whether this histone posttranslational modification is altered in the presymptomatic AD brain remains unknown.

To investigate changes in histone lactylation across different brain cell types in presymptomatic AD, we performed immunofluorescent co-staining of histone 4 lysine 12 lactylation (H4K12la) with antibodies targeting markers for neurons (NeuN), astrocytes (GFAP), and microglia (Iba1) in the hippocampus of 3-month-old 5XFAD mice. Most neurons exhibited H4K12la fluorescence **(Figure 2D),** with increased fluorescent intensity in neurons from 5XFAD mice compared to wild-type (WT) controls **(Figure 2E).** In contrast, we observed no significant differences in H4K12la intensity in astrocytes between 5XFAD and WT mice **(Figures 2F-G).** Additionally, no microglia were co-stained with the H4K12la antibody (Data not shown).

### Single cell transcriptional profiling of hippocampus in presymptomatic 5xFAD mice

To investigate the molecular mechanisms underlying altered BBB and cerebral metabolism in the presymptomatic 5xFAD mouse model, we performed single nucleus RNA-sequencing (snRNA-seq) of mouse hippocampal tissue from 3-month-old 5XFAD and WT control mice **(Figure S4A).** While vascular cells represent up to 10% of all brain cells, they are typically underrepresented by snRNA-seq of bulk tissue inputs^43^. We optimized protocols to enrich blood vessels based on published approaches^43^. Briefly, tissue was homogenized, and blood vessels were enriched by ultracentrifugation in a dextran gradient.

The solution was then passed over a mesh filter to capture blood vessels, which were subsequently eluted from the filter. This approach was optimized on the mouse cortex, resulting in a dramatic enrichment of blood vessels **(Figure S4B)** and vascular transcripts **(Figure S4C).** We then applied this approach to hippocampal tissue dissections, conducting both total nuclei preparation and brain vessel enrichment followed by nuclei preparation. After quality control, we annotated the transcriptomes of 75,175 nuclei **(Figure 3A),** encompassing eight excitatory neuron subtypes, inhibitory neurons, astrocytes, oligodendrocytes, oligodendrocyte precursor cells (OPCs), microglia, ependymal cells, and four vascular cell types (endothelial cells, pericytes, smooth muscle cells and fibroblasts). Endothelial cells could be further clustered into arteriovenous subtypes based on known marker genes (arterial, venous, capillary)^43,44^ **(Figure S5A).** We found that all vascular cell types displayed increased representation by snRNA-seq after blood vessel enrichment (p<O.05) **(Figure 3B-C).** For example, we observed that endothelial cells were 8-fold more abundant after vessel enrichment. Interestingly, we also found that glial cell types (astrocytes, oligodendrocytes, and microglia) were also increased after vessel enrichment. We next examined whether the cell type composition in the 5XFAD mouse model differed from that in wild type aged 3 months. We performed cell proportion analysis and found that there were no significant differences **(Figure 3D-E)** (p<0.05), suggesting that the observed functional vascular deficiencies are not due to a change in the number of vascular cells, which is consistent with our immunofluorescence-based measurements.

**Figure 3.**
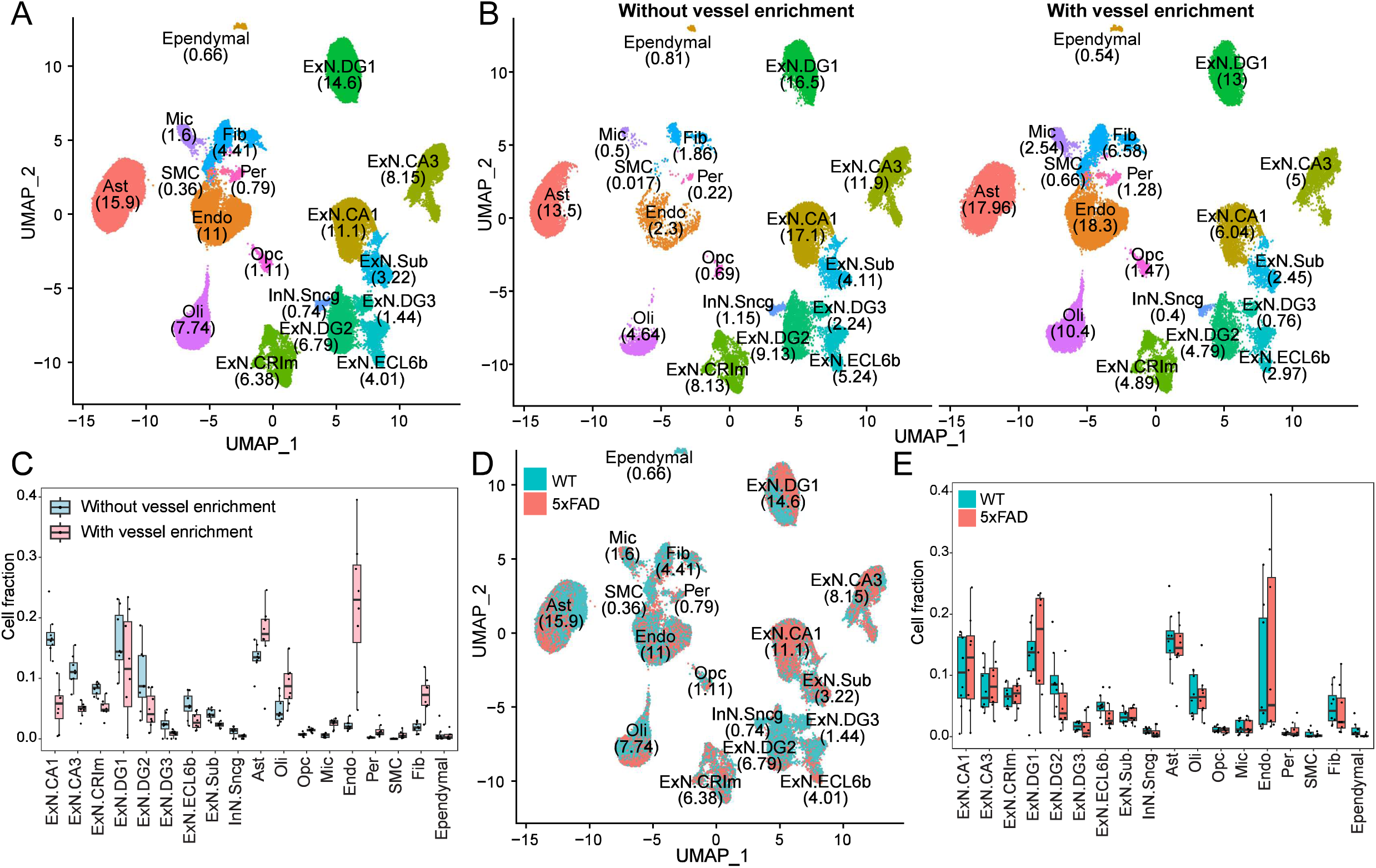
Transcriptional profiling of the hippocampus in presymptomatic 5xFAD mice at single-cell resolution. (A) UMAP (Uniform Manifold Approximation and Projection) plot visualizing 76,175 single nuclei annotated by cell type. The percentage of cells per cell type (%) is displayed. (B) UMAP visualization comparing conditions with (left) or without (right) blood vessel enrichment with cell type percentages included. (C) Cell proportion analysis indicates significant fractional differences in cell type proportions between conditions (with and without vessel enrichment). Statistical significance was determined using the Wilcoxon rank-sum test (n = 8 per condition, ‘adjusted p < 0.05). (D) UMAP visualization of all cells, colored to distinguish wild-type and 5XFAD mice. Cell type percentages (%) are shown for each genotype. (E) Cell proportion analysis highlights fractional differences in cell type proportions between wild-type and 5XFAD mice. Statistical significance was determined using the Wilcoxon rank-sum test (n = 4 per condition).

### Cell type-specific contribution to hypometabolism, impaired BBB integrity, and hippocampal acidification in 5xFAD mice

To evaluate the transcriptional alterations in hippocampal cell types of 3-month-old 5XFAD mice, we identified a total of 365 differentially expressed genes (DEGs), comprising 176 upregulated DEGs and 189 downregulated DEGs in 5XFAD mice **(Figure 4A, Supplementary Table 1).** Significant gene expression changes were observed in excitatory neurons within the CA1, DG1, and DG2 regions, as well as in astrocytes, oligodendrocytes, and vascular endothelial cells.

**Figure 4.**
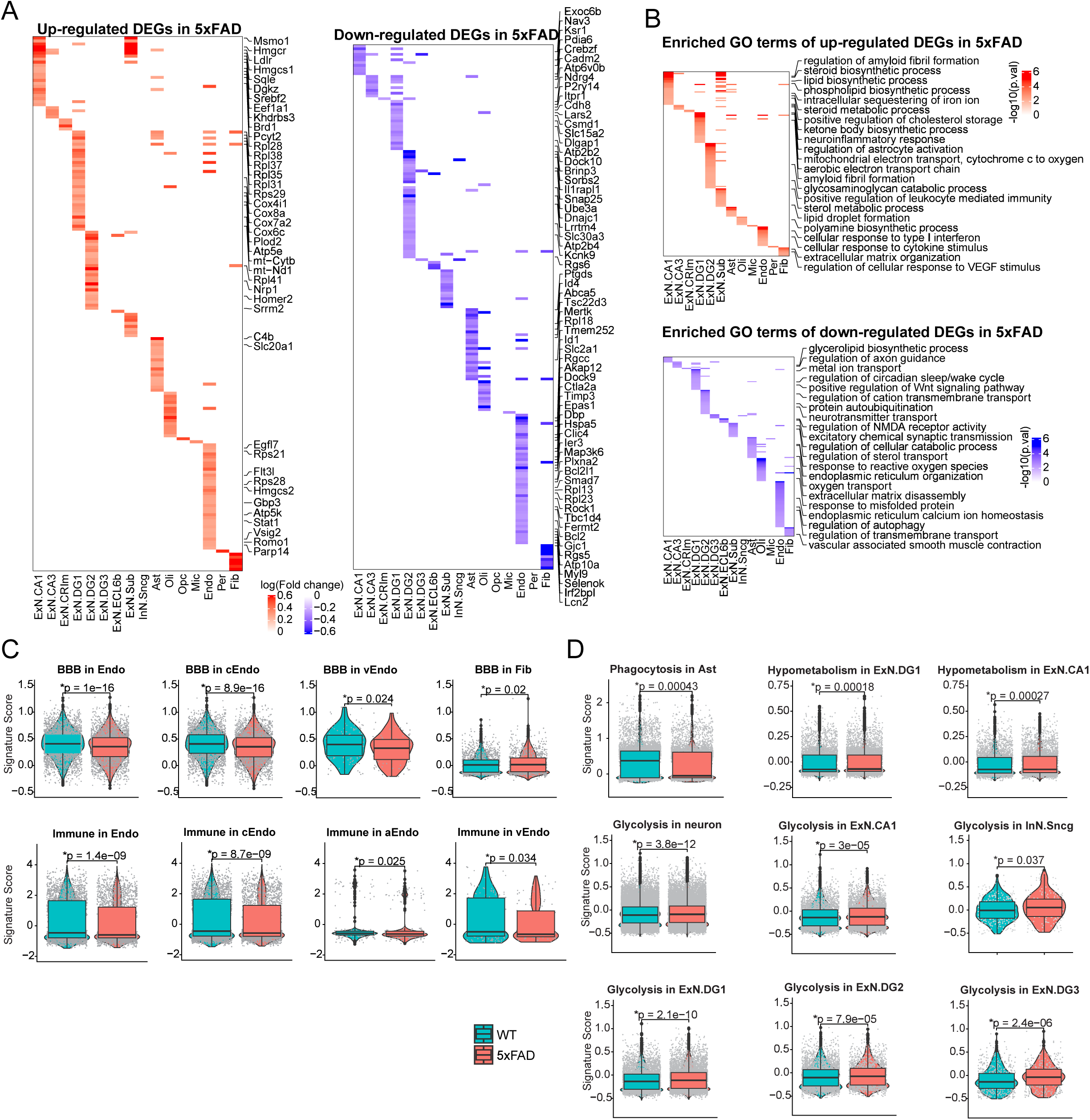
Transcriptional changes in hippocampal cell types in presymptomatic 5xFAD mice. (A) Heatmaps display significantly up-regulated (left) and down-regulated (right) genes across all cell types in 5XFAD mice. Values are presented as log-transformed fold changes, with representative top genes highlighted alongside the heatmaps. (B) Gene Ontology (GO) enrichment analysis of up-regulated and down-regulated genes in 5XFAD mice, categorized by cell type. Representative enriched GO biological processes are highlighted, with significance represented as —Iog10 of the p-value. A significance threshold of p < 0.01 was applied. (C-D) Gene signature score comparisons between wild-type (WT) and 5XFAD mice for selected biological functions in specific cell types, n = 4 mice/group. Violin plots illustrate the distribution of gene signature scores. Statistical significance was determined using the Student’s t-test (*p < 0.05).

We performed Gene Ontology (GO) enrichment analysis to elucidate the biological functions of DEGs across various cell types **(Figure 4B, Supplementary Table 2).** In the CA1 and DG1 excitatory neurons within the hippocampus of 5XFAD mice, genes involved in cholesterol biosynthesis and metabolism, such as *Msmol, Hmgcr, Hmgcsl, Sqle, Srebf2,* and *Pcyt2,* were significantly enriched among the upregulated genes **(Supplementary Table 3),** suggesting that lipid synthesis and homeostasis are disrupted in these neurons during the early stages of 5XFAD mice. As disruption in lipid homeostasis has been previously linked to AD progression, these results further implicate altered cholesterol metabolism in disease pathology.

In the hippocampus of 5XFAD mice, downregulated genes in CA1 and DG1 excitatory neurons were significantly enriched in functions related to lipid transport (e.g., *Exocβ),* axonal projection (e.g., *Nav3, Ndrg4),* and synaptic assembly, both presynaptic and postsynaptic (e.g., *Cadm2, Dlgapl, DockIO, Snap25, Sorbs2, Lrrłm4)* **(Supplementary Table 3). These** findings indicate an early disruption of synaptic transmission in the hippocampal excitatory neurons of 5XFAD mice, consistent with prior reports of early electrophysiological abnormalities in these mice ^45,46^. Furthermore, downregulation of genes associated with lipid transport and axonal projection suggests impairments in neuronal connectivity and communication.

Synaptic dysfunction is a hallmark of early AD, and alterations in genes like *Snap25* and *Dlgapl* have been implicated in synaptic vesicle cycling ^47,48^ and postsynaptic density organization^49,50^, respectively. These molecular changes may lead to cognitive deficits in the later stage of AD. Our findings align with previous research indicating that synaptic function is affected in the early stages of AD. These altered genes in the excitatory neurons may contribute to or be the consequence of the early occurrence of hypometabolism (reduced CMRO2) in the 5XFAD brain. Further targeting these genes and their involved pathways could clarify the therapeutic potential to mitigate neurodegeneration and cognitive decline in AD.

We also observed that genes responsible for immune response and inflammation *(Gbp3, Statľ)* were upregulated in the endothelial cells, while significantly downregulated genes in the endothelial cells included several involved in angiogenesis *(Dock9, Ctla2a, Plxna2, Dpb),* barrier function *(Askp12, Timp3)* and glucose transport (Slc2a1, Tbc1d4) **(Supplementary Table 2 and 3).**

To further dissect the molecular mechanism of cell-specific contribution to altered BBB permeability, hypometabolism, and acidification in the early stage of the 5XFAD mouse model, we selected genes that were known to be relevant with these functions **(Supplementary Table 2)** and calculated the signature scores for each cell, represented by the average expression of a group of genes for each functional term. We next compared the difference between 5XFAD and control cells and evaluated the statistical significance using a t-test **(Figure 4C-D).** We found genes relevant to BBB integrity and function are downregulated in endothelial cells, particularly in capillary endothelial cells (cEndo) and venous endothelial cells (vEndo) as well as in vascular fibroblasts **(Figure 4C, upper panel).** Genes involved in immune regulation and innate immunity were downregulated in the endothelial cells, including all types of endothelial cells **(Figure 4C, lower panel, Supplemental Tables 2 and 3).** These observations suggest that dysregulated genes in endothelial cells might be a major contributor to the early impairment of BBB integrity.

In addition, genes involved in phagocytosis were downregulated in astrocytes of 5XFAD hippocampus **(Figure 4D).** Genes leading to hypometabolism and glycolysis were mostly upregulated in excitatory neurons, including neurons in CA1, DG1, DG2, and DG3, as well as in inhibitory neurons **(Figure 4D).** These results suggest that reduced cerebral metabolism and increased acidification in the hippocampus of 5XFAD mice are associated with dysregulated genes in the neurons. Moreover, we confirmed that the downregulated gene *Slc2a1* **(Figure S5B)** was reflected in reductions in its protein product, glucose transporter 1 (Glutl), in hippocampal capillary vessels **(Figure S5C-D).**

### Exacerbated BBB leakage and amyloid β deposition in symptomatic 5XFAD mice

To determine whether the BBB becomes increasingly leaky as the disease model phenotype progresses, we examined BBB integrity in 9-month-old 5XFAD mice using EZ-link™ Sulfo-NHS-LC-Biotin leakage assessment, a marker for vascular permeability. Significant extravasation of biotin was observed in 5XFAD mice at this age **(Figure 5A-B),** indicating further compromised BBB function. This observation aligns with prior reports of BBB disruption in AD models ^9,13–26^. Additionally, WEPCAST MRI imaging and subsequent quantification revealed a statistically significant increase in BBB permeability in 5XFAD mice **(Figure 5C-D),** corroborating the histological findings.

**Figure 5.**
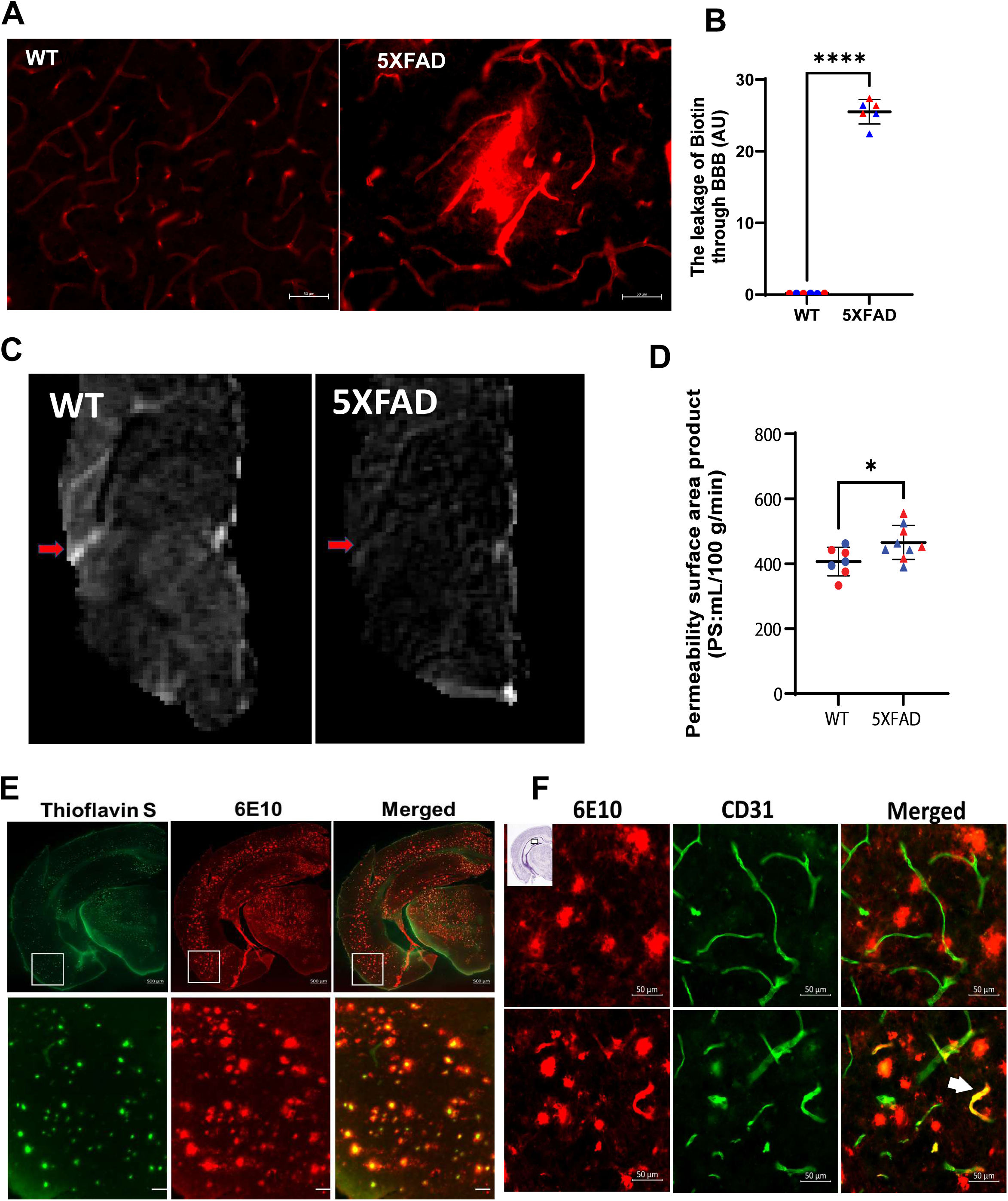
Exacerbated BBB leakage and amyloid β deposition in symptomatic 5XFAD mice. (A) Representative images show leakage of EZ-link™ Sulfo-NHS-LC-Biotin through BBB in the hippocampus of a 9-month-old 5XFAD mouse. Scale bar = 50 µm. (B) Quantification of biotin leakage into parenchyma in the hippocampus, n = 6 mice/ group, 3 male and 3 female. Red dots represent data from female mice, while blue dots represent male mice. ****p < 0.0001 by one-tailed student’s t-test. (C) Representative WEPCAST MRI images. (D) Quantification of permeability surface area product (PS) reveals significant increases in BBB permeability in 5XFAD mice, n = 7 WT (3 male and 4 female) mice and 9 AD (5 male and 4 female) mice. *p < 0.05, one-tailed Student’s t-test. (E) Brain-wide amyloid β deposition is labeled with Thioflavin S or anti-Aβ antibody (6E10) in a representative 9-month-old 5XFAD mouse. Scale bars: 500 µm (upper panel) and 10 µm (lower panel). (F) Amyloid β deposits are evident in cerebral blood vessels (arrow pointed) in a representative 9-month-old 5XFAD mouse. Scale bar = 50 µm.

Histological analysis revealed extensive Aβ pathology in 5XFAD mice at this age. Thioflavin S staining highlighted widespread amyloid fibrils, while immunolabeling with the 6E10 antibody confirmed the presence of Aβ plaques **(Figure 5E).** These findings are consistent with previous studies demonstrating aggressive Aβ accumulation in 5XFAD mice ^21^.

Furthermore, Aβ deposits were detected to colocalize with a subset of parenchyma microvessels in 5XFAD mice **(Figure 5F, arrow pointed)** or surrounding vessels **(Figure 5F),** consistent with the reported vascular pathology in this mouse model^51^, though we would mention that our results are in agreement with the previous report that only a limited number of microvessels show Aβ deposition. Our findings demonstrate that BBB leakage progressively worsened in this model, and our WEPCAST MRI is more sensitive in detecting early-stage, subtle BBB leakage (to small molecules like water). The progression of BBB leakage in this model may also explain some discrepancies among different studies, as different molecular weight molecular probes and different age groups were studied ^27^.

## Discussion

The motivation for this study was to understand early pathogenesis driving neurodegeneration and cognitive decline and identify potentially reversible biomarkers to facilitate the development of therapies for AD. We employed multiple advanced MRI techniques applicable to human patients, combined with snRNA-seq, to examine the 3-month-old 5XFAD mouse brain. First, we demonstrated that alterations in BBB permeability precede significant Aβ deposition and cognitive impairment. Notably, the increased permeability to water molecules without substantial extravasation of larger tracers suggests the sensitivity of WEPCAST MRI in detecting subtle BBB disruptions. Examination of neurovascular unit components revealed selective astrocyte activation, while tight junction integrity, pericyte markers, and endothelial cells remained unchanged, suggesting that the observed BBB changes may be reversible. Additionally, we identified significant reductions in CMRO_2_, indicating early metabolic disruption. Regional brain acidosis was observed in the hippocampus, as evidenced by CrCEST MRI, correlating with increased neuronal histone lactylation, suggesting metabolic shifts that may influence epigenetic regulation. Lastly, snRNA-seq analysis revealed altered gene expression in endothelial cells and hippocampal excitatory neurons, implicating BBB dysfunction and synaptic deficits in early disease progression. By 9 months, 5XFAD mice exhibited significant Aβ accumulation and BBB leakage to larger molecules, confirming the progression to advanced AD pathology. Given the significant BBB leakage and brain-wide Aβ accumulation observed in 9-month-old 5XFAD mice, our results reinforce the importance of targeting BBB preservation, metabolic restoration, and synaptic protection in efforts to delay or prevent AD progression.

The integrity of the BBB is crucial for maintaining central nervous system homeostasis. Using WEPCAST MRI, we detected increased BBB permeability to water molecules in 3-month-old 5XFAD mice. It is worth mentioning that WEPCAST MRI quantifies the relative fraction of magnetically labeled water spins that penetrate brain tissue versus those remaining in veins. Specifically, water spins from the blood can enter the brain tissue through the BBB. Increased water spins in arterial blood indicate “leakage” into brain tissue upon BBB disruption. Quantitatively, the fraction of water spins entering brain tissue is termed the water extraction fraction. The WEPCAST MRI benefits from high sensitivity to subtle BBB leakage by employing small water molecules (18 Da) as tracers. Using this technique, we previously found that patients with mild cognitive impairment (MCI, an early form of AD) manifested an increased BBB permeability to water, and BBB permeability to water is significantly associated with CSF Aβ42/Aβ40 ratio ^26^. The current findings suggest that the 5XFAD model recapitulates subtle BBB breakdown in the presymptomatic stage.

Histological analysis using EZ-link™ Sulfo-NHS-LC-Biotin did not detect significant extravasation in 3-month-old 5XFAD mice, suggesting that early BBB disruption permits passage primarily of small molecules, such as water. Additional studies corroborate that many aspects of BBB integrity are preserved in 5XFAD mice ^27^. Our observations suggest a subtle increase in BBB permeability precedes substantial Aβ deposition, as only minimal Thioflavin S-positive Aβ fibrils were observed in the frontal cortex and subiculum at this age. Cognitively, these mice performed comparably to wild-type controls in Y-maze and novel object recognition tests, indicating preserved cognitive function despite the subtle BBB alterations.

Further histological examination of the neurovascular unit components in 3-month-old 5XFAD mice revealed mild gliosis, including increased GFAP and microglia exhibited activation morphology. However, tight junction protein claudin-5, pericyte, endothelial cell, and aquaporin-4 polarization remained unchanged. These findings suggest changes in BBB permeability at this early stage are most likely reversible. Our results underscore the importance of targeting BBB preservation in therapeutic strategies aimed at delaying or preventing disease progression.

Moreover, the brain has high oxygen demand, which necessitates precise regulation of oxygen delivery to maintain proper neuronal function. In AD conditions, regional hypoperfusion and decreased tissue oxygenation have been documented^52–54^, correlating with cognitive decline and neurodegeneration. Our findings of significantly reduced CMRO and OEF in 3-month-old 5XFAD mice suggest that metabolic impairments occur much earlier in the disease progression. These reductions may lead to neuronal dysfunction and exacerbate Aβ deposition.

Extracellular acidosis has been observed in postmortem AD brains, and brain tissue acidification can enhance amyloid plaque deposition^36,38^. Our CrCEST MRI measures revealed significantly lower pH levels in the hippocampus of 3-month-old 5XFAD mice compared to controls, but not other brain regions. This region-specific acidification may serve as an early biomarker for AD; it may imply underlying neuroinflammatory processes. Moreover, histone lactylation, a post-translational modification influenced by increased lactate in the local environment, has been implicated in regulating gene expression. Increased H4K12 lactylation has been reported in symptomatic 5XFAD mice and postmortem AD brains^42^. We demonstrated elevated H4K12la in the hippocampal neurons of 3-month-old 5XFAD mice, without changes in astrocytes or microglia, implying a possible metabolic shift from aerobic to anaerobic metabolism in neurons of presymptomatic 5XFAD mice. These metabolic changes may contribute to epigenetic modifications and consequently lead to AD pathology.

The snRNA-seq studies have provided insights into cell type-specific molecular changes in the hippocampus of presymptomatic 5XFAD mice, particularly concerning BBB function and cerebral metabolic changes. The compromise in BBB integrity is associated with altered expression of genes involved in endothelial cell function, particularly capillary endothelial cells. Defining these transcriptional changes highlights novel molecular targets for future therapeutic validation.

Synaptic deficits are a hallmark of AD, and our snRNA-seq analyses have revealed alterations in the gene expression critically involved in synaptic function in presymptomatic 5XFAD mice, consistent with other reports that synaptic changes are early events in human AD^55–57^. Notably, genes essential for synaptic vesicle transport and connectivity are found to be downregulated in the hippocampal excitatory neurons of presymptomatic 5XFAD mice.

These findings offer insights into the pathophysiological mechanisms underlying early AD and suggest potential biomarkers and therapeutic targets for early intervention. Future studies should explore whether interventions aimed at stabilizing BBB integrity and restoring metabolic balance in presymptomatic stages can mitigate disease onset and progression.

## Resource Availability

### Lead Contact

Requests for further information and resources should be directed to and will be fulfilled by the lead contact, Wenzhen Duan.

### Material Availability

This study did not generate new unique reagents.

### Data Availability

Data and code availability

The single-cell RNA-seq dataset generated in this study will be made publicly available via the Gene Expression Omnibus under accession number GSE295077 upon publication. Additional supporting datasets are available from the corresponding author upon reasonable request. All code used for data processing and analysis will be released at https://github.com/nasunmit/5xfad at the time of publication. Microscopy data and MRI data reported in this paper will be shared by the lead contact upon request. Any additional information required to reanalyze the data reported in this paper is available from the lead contact upon request.

## Acknowledgements

This project is supported by R01NS124084 and R01NS127344 (to W.D.), R01NS129032 and U01DA053631 (to M.H. and M.K.), R01AG080104 (to J.X.), R01AG081932 (toZ.W.), R01AG071515 and P41EB031771 (to H.L.). R.M.L. was supported by a postdoctoral fellowship from NIH/NINDS (F32NS128067). High-throughput sequencing was performed at The Single Cell & Transcriptomics Core located at the Johns Hopkins School of Medicine.

## Author Contributions

MY and WD conceptualized the study, designed overall experiments, analyzed/interpreted results, and wrote the manuscript. NS analyzed snRNA-sequencing data analysis and wrote the manuscript. RL optimized and performed blood vessel enrichment, conducted snRNA-seq, and wrote the manuscript. ZW conducted WEPCAST and TRUST MRI scans, as well as data analysis. AK, YO, YJ, RL conducted immunohistology experiments and image analysis. HL designed the physiological MRI sequences and conceptualized the MRI BBB and CMRO2 measurements in AD mice. ZZ, AL, JX conducted CrCEST MRI experiments and image analysis. LD, YZ, HW conducted EZ-link™ Sulfo-NHS-LC-Biotin BBB leakage experiments and analyzed data. MK provided mentorship to NS for snRNA-seq analysis. MH supervised RL and co-wrote the manuscript.

## Supplemental Information

Supplementary Table 1. All DGEs

Supplementary Table 2. DGEs and Goenrichment

Supplementary Table 3. Dysregulated genes and their functions

Supplementary Table 4. DGEs in bulk vs Vascular enriched samples Pseudobulk results

Supplementary Table 5. Mean and SD for all figures

Supplementary Table 6. Antibody information

## STAR Methods

### Experimental Model and Subject Details

#### Animals

5XFAD transgenic line (B6.Cg-Tg(APPSwFILon,PSEN1*M146L*L286V)6799Vas/Mmjax/J stock no. 034848) was obtained from the Jackson Laboratory and bred in our lab. Mice were genotyped at 4 weeks of age via tail biopsy using a fee-based service provided by Transnetyx Inc. Both male and female mice, as well as littermate control wild type (WT) mice, were used, and individual data points are presented in all graphs. Mice were housed in sex-matched groups of five in standard mouse cages on a 12-h light/dark reversed cycle at a room temperature (RT) of 23 °C, with free access to food and water. All procedures were approved by the Animal Care and Use Committee of the Johns Hopkins University.

#### MRI measures

Functional and physiological MRI were conducted at an 11,7T Bruker Biospec system (Bruker, Ettlingen, Germany) with a horizontal bore equipped with an actively shielded pulse field gradient (maximum intensity of 0.74 T/m). Images were acquired using a 72-mm quadrature volume resonator as a transmitter, and a four-element (2×2) phased-array coil as a receiver. The homogeneity of the BO field over the mouse brain was optimized with a global shimming (up to 2nd order) based on a subject-specific pre-acquired field map.

Functional MR imaging was conducted under low-dose isoflurane anesthesia carried by medical air (21% 02, 78% N2). Anesthesia induction was at 1.5% concentration for 15 minutes, followed by maintenance at 1.0%. The respiration rate of each mouse was continuously monitored during experiments to ensure survival and maintain consistent respiratory rates across all experimental mice. If a mouse exhibited a breathing rate exceeding 150 breaths per minute, the maintenance isoflurane dose was slightly increased to 1.2%. This anesthesia protocol has been previously utilized and documented ^58^. Additionally, each mouse was immobilized using a bite bar and ear pins, and then placed on a water-heated animal bed with temperature control.

***Oxygen extraction fraction (OEF)*** is defined as the arteriovenous difference in blood oxygenation and was measured using the T2-relaxation-under-spin-tagging (TRUST) MRI technique. TRUST scan was implemented following the reported MRI protocol^31^. Key parameters were: TR/TE = 3500/6.5 ms, FOV = 16×16 mm^2^, matrix size = 128×128, slice thickness = 0.5 mm, inversion-slab thickness = 2.5 mm, post­labeling delay = 1000 ms, eTE = 0.25, 20, 40 ms, and scan duration = 5.6 min with two repetitions.

Cerebral blood flow (CBF) was evaluated with phase contrast (PC) MRI focusing on the three major feeding arteries of the brain (left/right internal carotid arteries and basilar artery). The previously reported protocol was utilized^53^, and the key parameters were: TR/TE = 15/3.2 ms, FOV = 15*15 mm^2^, matrix size = 300*300, slice thickness = 0.5 mm, receiver bandwidth = 100 kHz, flip angle = 25°, and scan duration = 0.4 min per artery. Brain volume was estimated from a T2-weighted fast-spin-echo MRI protocol (TR/TE = 4000/26 ms, FOV = 15×15 mm^2^, matrix size = 128*128, slice thickness = 0.5 mm, 35 axial slices, and scan duration = 1.1 min)^31^.

***Blood-brain barrier*** function was assessed with water extraction with phase-contrast arterial spin tagging (WEPCAST) MRI^5^. Key parameters were: TR/TE = 3000/11.8 ms, labeling duration = 1200 ms, FOV = 15*15 mm^2^, matrix size = 96×96, slice thickness = 1 mm, labeling-pulse width = 0.4 ms, inter­labeling-pulse delay = 0.8 ms, flip angle of labeling pulse = 40°, and scan duration = 4.0 min with two-segment spin-echo echo-planar-imaging acquisition.

#### Creatine-CEST (CrCEST)

The CEST experiments were performed using continuous-wave CEST (cwCEST). MR images were acquired using a Turbo Spin Echo (TSE) sequence with TE = 18 ms, TR=5 s, TSE factor = 20, a matrix size of 64*64. The slice thickness was 1.5 mm and the FOV was 16 *16 mm^2^. The saturation field strength (B-i) and length for CrCEST were 2 µT and 1 s, respectively, according to previous studies ^60,61^. The frequency range measured to assess CrCEST was 1.00 to 3.50 ppm with an increment of 0.05 ppm, and the one for amideCEST was 2.30 ppm to 5.00 ppm with an increment of 0.10 ppm. SO images for the CEST studies were acquired by setting the offset at 200 ppm. ATi map was acquired using the RAREVTR sequence (RARE with variable TR =0.5, 1,1.5, 2, 3.5, 5, and 8 s). All CEST MRI images were processed using custom-written MATLAB scripts (MathWorks, www.mathworks.com, version 9.8.0.1417392 (R2020a) Update 4). The extraction and quantification of the CrCEST signal were achieved using polynomial and Lorentzian line-shape fitting (PLOF) as detailed previously^61^.

#### Behavioral testing

***Y-maze*** was conducted at 3 months of age. Briefly, mice were exposed to the behavioral test room 2h before the formal test as habituation. Mazes were cleaned with 70% ethanol and allowed to dry before each test began. Then, each mouse was gently and randomly put into one arm of the Y-maze and allowed to explore for 10 min. The rodent was considered to have entered an arm when its whole body, minus its tail, was in the arm. All behavior performance was recorded by the camera above the Y-maze and analyzed by Anymaze software (Stoelting, USA). Percent of spontaneous alternation is defined as the number of spontaneous alternations divided by the total possible triads.

***Novel Object Recognition (NOR)*** was also conducted at 3 months of age. In brief, after habituation in the room for 2h, mice from each cohort were acclimatized to a 50 cm×50 cm testing chamber for 2 min. After acclimatization, the mice were removed, the testing area was cleaned with 70% ethanol, and two identical toy bricks were placed in the corners of the testing area, approximately 5 cm from each wall, for 10 min. The mice were removed, the testing area was cleaned with 70% ethanol, and one object was replaced with a novel toy brick with a different texture and shape. Mice were reintroduced, and the ratio of both the number of visits and the time spent at each object was recorded via Anymaze software (Stoelting, USA) for 10 min.

#### BBB permeability assay

Blood brain barrier permeability was assessed by using the EZ-Link™ Sulfo-NHS-biotin (Thermo Fisher, 21217, Mr. 556.59) tracer. Mice were injected with freshly made EZ-Link™ Sulfo-NHS-biotin solution (50 mg/ml) at 500 mg/kg body weight and perfused with 4% paraformaldehyde (PFA) in PBS after 30 min. Brains were carefully dissected, and 40 µm thick brain slices were incubated with Texas Red-conjugated streptavidin (Vector Lab, SA-5006-1) for 30 min, then washed for another 30 min. Brain sections were imaged using a Zeiss Microscope, and images of biotin tracer track were obtained. The methods for signal quantitative analysis are as follows: Hippocampal regions of brain parenchyma are randomly selected in each brain section, and the optical density value was measured by ImageJ software to reflect the signal strength of the leaked Sulfo-NHS-Biotin. 6 mice/genotype (3 male and 3 female) were sectioned, and three brain slides of each sample were analyzed.

#### Thioflavin S (TS) staining

Thioflavin S (Sigma Aldrich, T1892-25G, TS) was used to visualize amyloid deposits and was prepared according to the manufacturer’s instructions. Tissue slices were dehydrated sequentially in 75%, 85%, and 90% ethanol, then incubated with 0.002% TS in the dark for 10 min. They were then rehydrated in 90%, 85% and 75% ethanol and washed with 1x PBS three times, each for 10 min. Afterward, brain sections were mounted for imaging or processed for further immunostaining with other antibodies indicated in the study.

#### Western blotting

Freshly frozen brain tissues with dissected brain regions were lysed in RIPA lysis buffer containing a cocktail of protease and phosphatase inhibitors (Cat # 5872, Cell Signaling Technology). The protein concentration of lysates was measured by BCA assay and was adjusted to the final concentration of 1 µg/µl. After heat denaturation, equal amounts of protein in the lysates were separated by SDS-PAGE and then transferred to a polyvinylidene fluoride (PVDF, Millipore) membrane. The membranes were blocked with 5% non-fat milk in tris-buffered saline containing 0.1% Tween-20 (TBST) for 1 h at room temperature followed by incubation with the indicated primary antibodies overnight at 4°C. After three washes with TBST, the membranes were incubated with horseradish peroxidase (HRP)­conjugated secondary antibodies. The immunoreactive proteins were then detected using an enhanced chemiluminescent (ECL) substrate and visualized with the imaging system (Odyssey). For antibody details, see Supplementary Table 6. The band intensities were quantified with ImageJ software.

#### Immunohistochemistry and quantification

Mice were anesthetized with isoflurane and sequentially perfused transcardially with phosphate-buffered saline (PBS) and 4% paraformaldehyde in PBS. Fixed and sucrose-cryoprotected brain tissue was sectioned coronally with a cryostat (Leica, CM1950) into 40-µm sections for immunohistochemistry. Staining was performed on free-floating mouse brain sections. The sections were rinsed three times for 10 min in PBS and then incubated with blocking buffer (5% goat or donkey serum and 0.2% Triton X-100 in PBS) for 1 h at room temperature. The sections were then incubated with primary antibodies (details in Supplementary Table 6) in blocking buffer overnight at 4 °C. After thorough washes in PBS, sections were incubated with a 1:1,000 dilution of Alexa 488-, Alexa 555-, and Alexa 647-conjugated secondary antibodies (Thermo Scientific, Goat anti-rabbit 488: A11008, Goat anti-mouse 488: A11001, Goat anti­rabbit 555: A32732, Goat anti-mouse 555: A21422, Goat anti-rat 647: A21247) appropriate for the species of the primary antibodies at room temperature for 1 h, followed by Hoechst counterstains. The sections were examined, and images were acquired using Zeiss LSM700 laser-scanning conſocal microscopes. For quantitation of cell lactylation and fluorescence intensities of IBA1, GFAP, and Glutl in the hippocampal sections, images of brain regional areas were acquired using a Zeiss LSM700 microscope with a × 20 objective. Analyses were performed using Fiji ImageJ software. For quantification, five consecutive sections per mouse were used, with five mice in each group. From each section, four images were selected for quantification. Sum intensity values were calculated from these images to obtain results for statistical analysis.

#### Tissue dissection

Mice were anesthetized with isoflurane and rapidly decapitated. Brains were harvested and dissected immediately on an ice-dry ice mixture. Hippocampal dissections were flash-frozen in liquid nitrogen and stored at −80°C prior to further processing.

#### Vascular and non-vascular nuclei preparations

Vascular enrichment protocols were adapted from those previously described ^62^. The following solutions were prepared fresh: (1) homogenization media: 0.5% bovine serum album (Millipore Sigma, #126609) in MCDB131 media, (2) dextran solution (18.75% dextran in 1xPBS), (3) hypotonic solution (20 mM Tris-HCI, 10 mM NaCI, 3 mM MgCI2, 1 mM DTT, with pH of 7.4 prepared in nuclease-free water), and (4) nuclei buffer (0.5% BSA and 1 mM DTT in 1xPBS). Each was supplemented with 5 µL mL-1 SUPERase·ln™ RNase Inhibitor (ThemnoFisher #AM2696) and 10 µL mL-1 RNasin® Ribonuclease Inhibitor (Promega #N2515) and kept on ice during use. The tissue was transferred to a prechilled 2 mL dounce homogenizer containing 1 mL of homogenization media and homogenized for 10 strokes per pestle, loose and tight. Homogenate was then diluted with an additional 1 mL of homogenization media and split across two Dolphin microcentrifuge tubes, one tube used for vascular nuclei preparations with 1.8 mL volume (90% of tissue), and one tube used for standard nuclei preparations using 200 µL volume (10% of tissue). Tissue was pelleted by centrifugation at 2000 x g for 10 min in a swinging bucket rotor. For vascular nuclei preparations, pellets were resuspended into 2 mL of dextran solution and centrifuged at 14,000 rpm for 15 min in a refrigerated Optima MAX-TL Ultracentrifuge (Beckman Coulter, #A95761). The supernatant containing a myelin layer was removed and the pellet was resuspended into 0.5 mL of hypotonic solution. Microvessels were then captured by gently pipetting the resuspended vascular-enriched pellet onto a 20 µm pluriStrainer (pluriSelect, #435-50020-01). 1 mL of hypotonic solution was washed over the filter to further deplete non-vascular cells. Then microvessels were eluted from the filter by forceful pipetting of 1.4 mL of hypotonic solution after flipping the filter over a new conical tube. Solutions containing microvessels were incubated on a shaker for 10 min and then supplemented with 0.15% NP-40 (BioProcessing Biochemicals, #25,030) and 15 mM DHPC (Avanti #850306P). After pipette mixing, solutions were incubated for 30 min and then transferred to a 4 mL Potter-Elvehjem tissue grinder (DWK, #885510-0020) with mechanical disruption conducted at 900 rpm for 25 strokes with the associated PTFE pestle. Resulting solutions were collected and filtered from a FlowMi 40 µm strainer, supplemented with 0.16 mLof 5% BSA, vortexed at full speed for 10 sec to homogenize nuclei, and spun down at 550 x g for 10 min. For non-vascular nuclei preparations, previously reported methods were used^63^

#### Quantitative PCR validation of vascular enrichment

Paired vascular and non-vascular nuclei preparations from flash-frozen mouse cortex were inputted into the Single Cell-to-CT™ qRT-PCR Kit (Invitrogen™, #4458236) following standard manufacturer protocols. TaqMan primers for *Ipo8* (housekeeping gene), *Cldn5, Abcbla, Pecaml, Slc2a1, Mfsfd2a* (endothelial-specific), *Acta2, Pdgfrb* (mural-specific) and *Aqp4, AldhM, Mog, Syp, Rbfox3, Nefh* (non-vascular) transcripts were used. We repeated this experiment across two biological replicates of the mouse hippocampus, finding ∼24-fold enrichment of *Cldn5,* and ∼2-fold enrichment of *Rbfox3* using the delta-delta Ct method of quantification.

#### Single nucleus RNA sequencing (snRNA-seq)

Paired vascular and non-vascular nuclei preparations were loaded onto a Chromium Next GEM Chip G (lOxGenomics, PN-1000120) with libraries prepared using a Chromium Next GEM Single Cell 3’ Kit v3.1 following manufacturer recommendations (lOxGenomics, PN-1000269). In total, four 5XFAD and four WT mice were profiled, corresponding to 16 total samples (bulk and vascular for each). For each sample, we targeted 6,000 nuclei on average. Libraries were sequenced on a NovaSeq S4 (lllumina) targeting 60,000 reads per nucleus. We obtained 59,058 reads per cell on average.

#### snRNA-seq data analysis

We aligned the raw reads to the mouse genome and estimated the gene counts using the CellRanger software (v3.0) (10x Genomics). We then used the Seurat (v4.3.0.1) package in R for downstream analysis. We selected cells with more than 500 detected unique molecular identifiers (UMIs) from protein­coding genes for further analysis. We also use the ratio of mitochondrial genes to measure the quality of cells (cells with higher than 5% were removed). We used DoubletFinder with by default parameters to remove the potential doublets and DecontX to remove the ambient RNA contamination (https://genomebiology.biomedcentral.eom/articles/10.1186/s13059-020-1950-6) from snRNA-seq data. We used the top 2000 highly variable genes for Principal Component Analysis (PCA). The first 30 PCs were used for non-linear dimensionality reduction (UMAP) for 2-D visualization. The “FindMarkers” function in Seurat was used to identify marker genes for each cluster and cell type using the default parameters. We also used the FindMarkers function to identify differentially expressed genes (DEGs) between 5XFAD and control mice, employing the MAST method while considering covariates, including contamination score, percentage of mitochondrial genes, and ribosomal genes. To annotate vascular cells with high resolution, we used previous datasets enriched for vascular cell types to recognize subtypes of vascular cells^43^.

#### Cell fraction analysis

We evaluated the statistical significance for compositional differences between phenotypic groups of interest using the Wilcoxon rank-sum test (AD groups, w/wo vessel enrichment). We used an adjusted p-value of less than or equal to 0.05 as the cutoff for significance.

#### Gene Ontology enrichment analysis

We used Enrichr in R to perform the enrichment analysis for Gene Ontology Biological Processes (Benjamini-Hochberg adjusted p-value < 0.05 as a cutoff for DEGs) and selected the representative terms of interest for visualization using a heatmap.

#### Gene signature analysis

*We* defined a series of functional terms that included BBB, immune, phagocytosis, hypometabolism, and glycolysis by manually selecting genes with known functions in each term (Supplementary table). We next calculated the signature scores (i.e., BBB, immune, phagocytosis, etc) for each cell, which were represented by the average expression of a group of genes for each functional term. We evaluated the statistical significance of differences between 5XFAD and control cells using the t-test.

#### Statistical Analysis

Data are expressed as individual values unless otherwise noted. Statistical analysis was performed with SPSS using two-tailed Student’s t-test, one-way ANOVA and two-way ANOVA with Bonferroni post-hoc tests. The P-values less than 0.05 were considered statistically significant. N is reported in the figure legends.

## Supplemental Text and Figures

**Figure S1.**
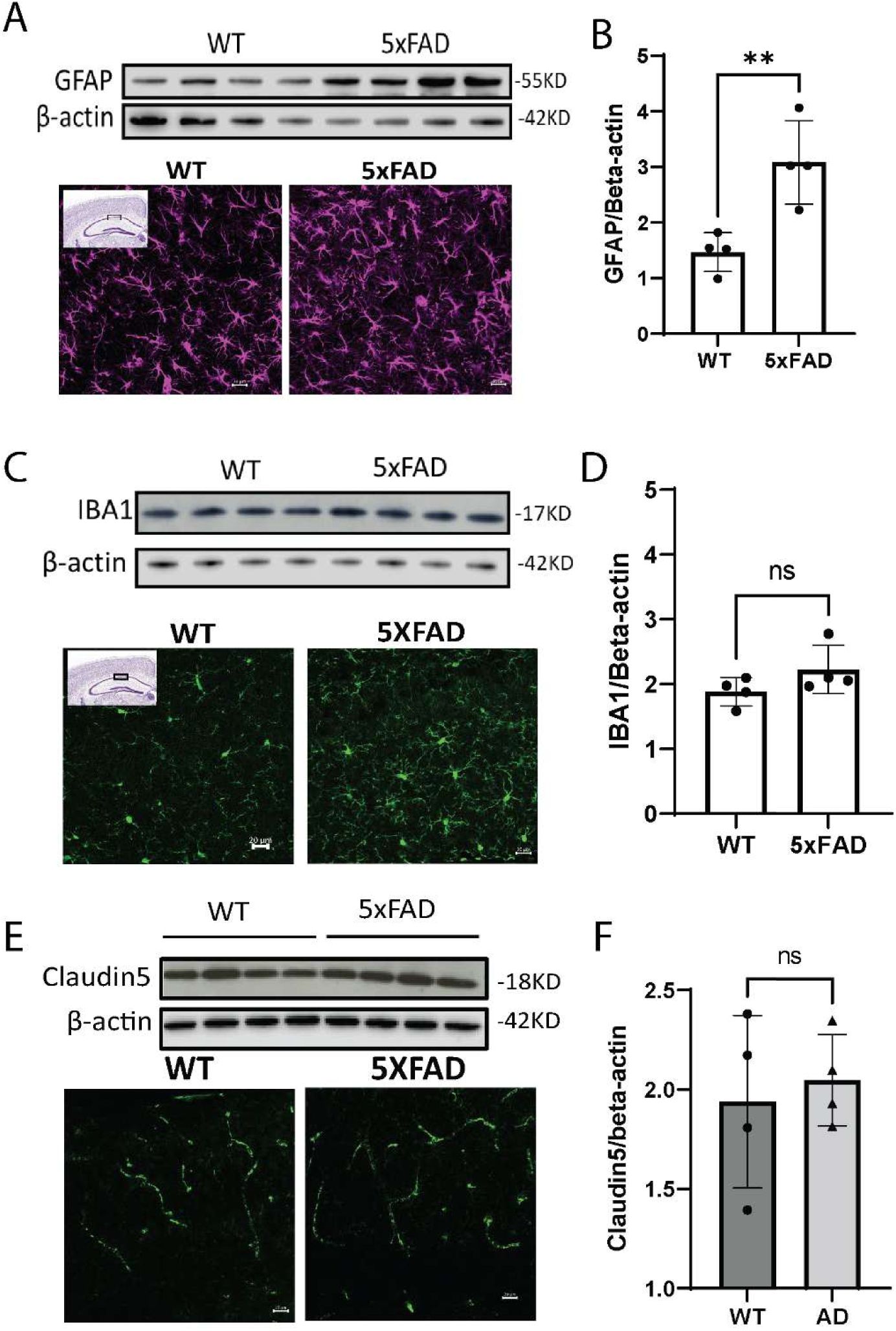
Gliosis in prodromal 5XFAD mice. (A) GFAP Western blots and representative immunofluorescent images reveal astrogliosis in the hippocampus of prodromal 5XFAD. (B) Quantification of GFAP protein levels from Western blots shows a significant increase in 5XFAD mice compared to controls, n=4. (**p < 0.01, Student’s t-test). (C) Western blots and immunofluorescent staining of the microglial marker IBA1 indicate activated microglia morphology in 5xFAD mice. (D) Quantification of IBA1 protein levels between AD and control mice. n=4. (E) Western blot analysis and immunofluorescent staining of the tight junction protein Claudin 5 indicate alterations in BBB integrity in 5XFAD mice. (F) Quantification of Claudin 5 protein levels from the Western blots. n=4. Statistical analysis confirms no significant changes (ns, Student’s t-test).

**Figure S2.**
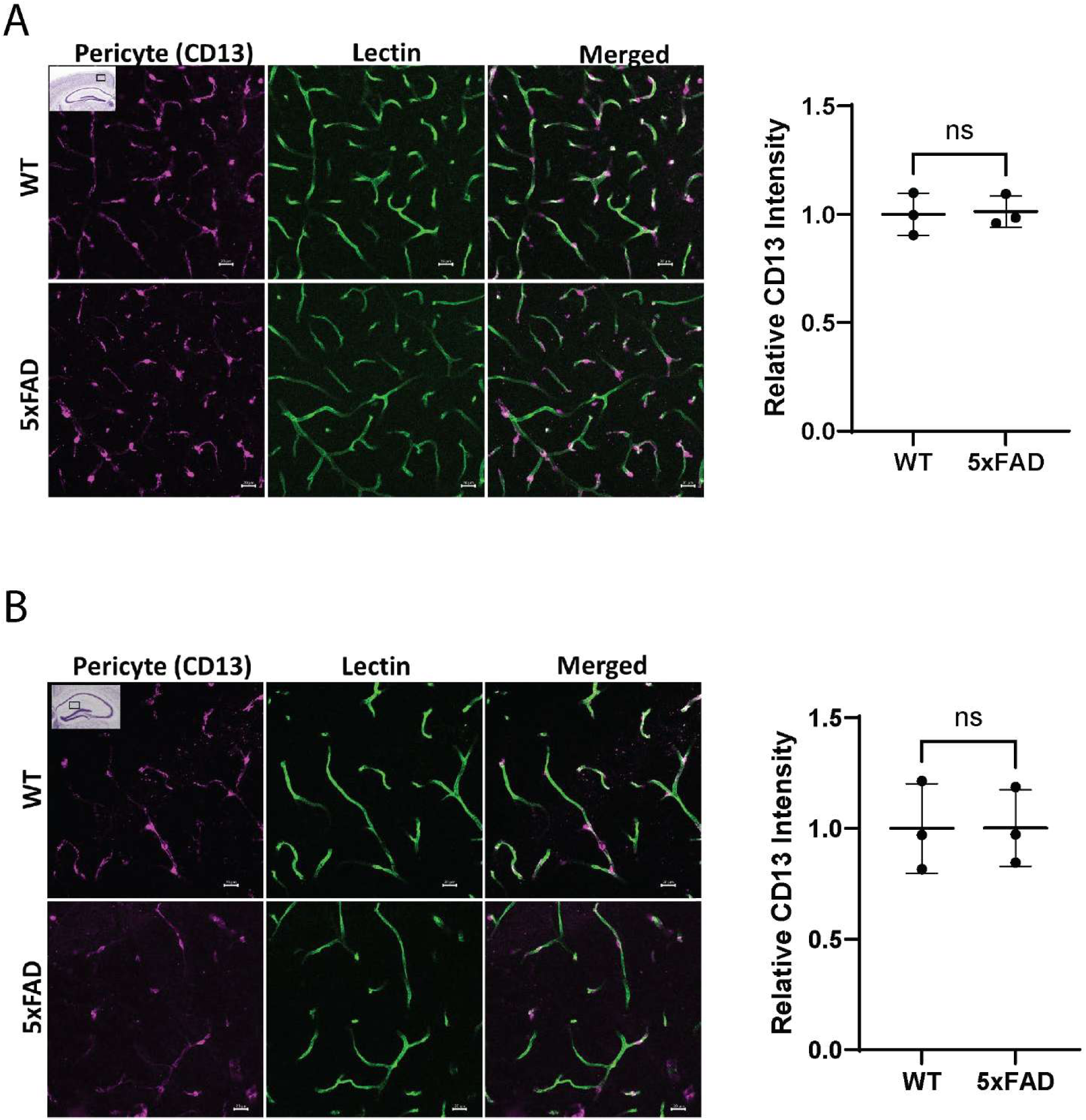
Preservation of pericyte intensity in prodromal 5XFAD mice. (A) Representative immunofluorescent staining of the pericyte marker protein CD13, alongside the vascular marker lectin, shows no significant difference in CD13 fluorescence intensity in the cerebral cortex of 5XFAD mice compared to controls. n=3. (B) Representative immunofluorescent staining and quantification of CD13 fluorescence intensity in the hippocampus show no significant difference between the two groups. n=3. Statistical analysis confirms no significant changes (ns, Student’s t-test).

**Figure S3.**
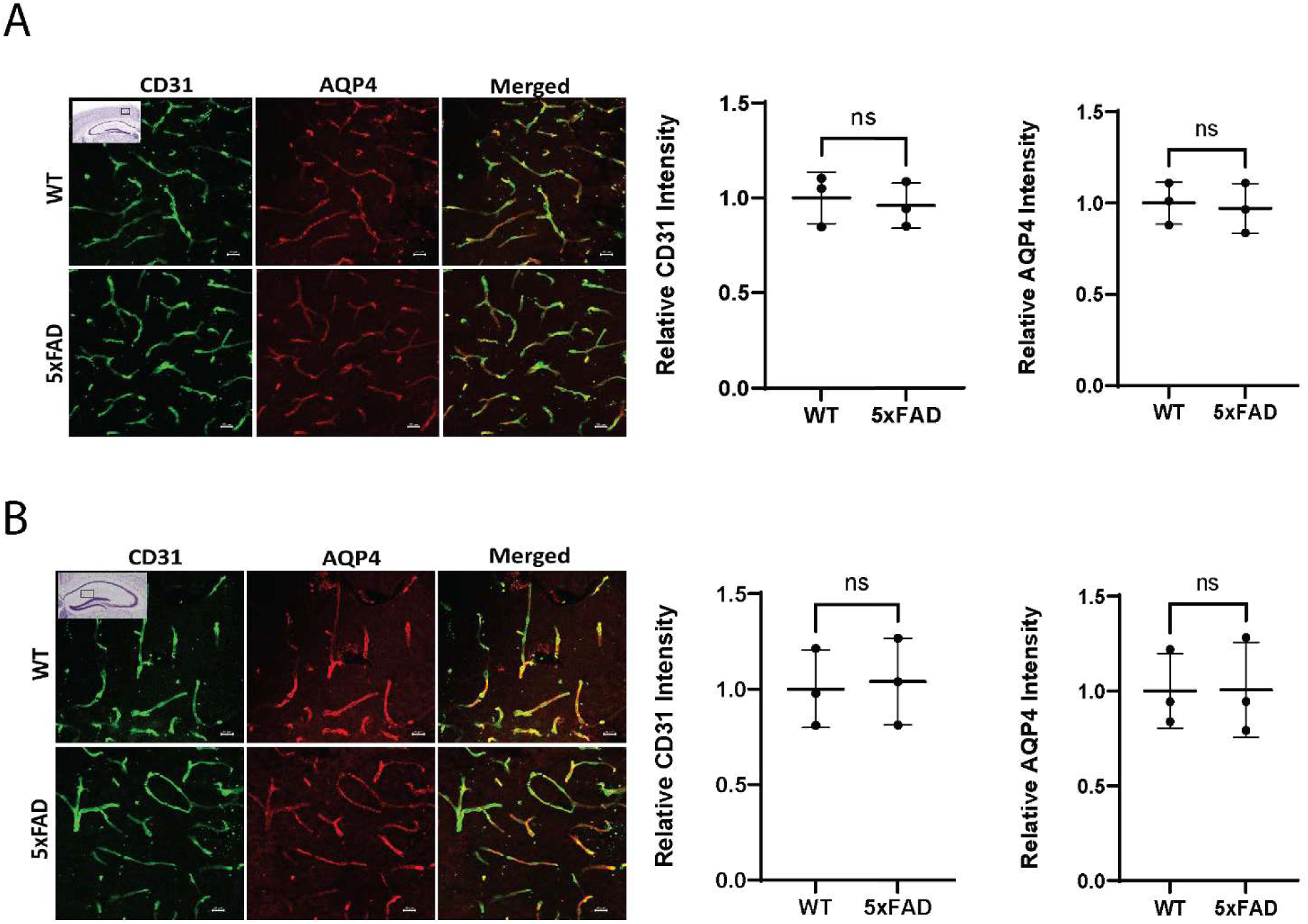
Preservation of endothelial cell intensity and AQP4 polarization in prodromal 5XFAD mice. (A) Representative immunofluorescent staining of the endothelial marker protein CD31 and the water channel protein AQP4 shows no significant difference in fluorescence intensity in the cerebral cortex of 5XFAD mice compared to controls. n=3. (B) Representative immunofluorescent staining and quantification of CD31 and AQP4 fluorescence intensity in the hippocampus show no significant difference between groups. n=3. Statistical analysis confirms no significant changes (ns, Student’s t-test).

**Figure S4.**
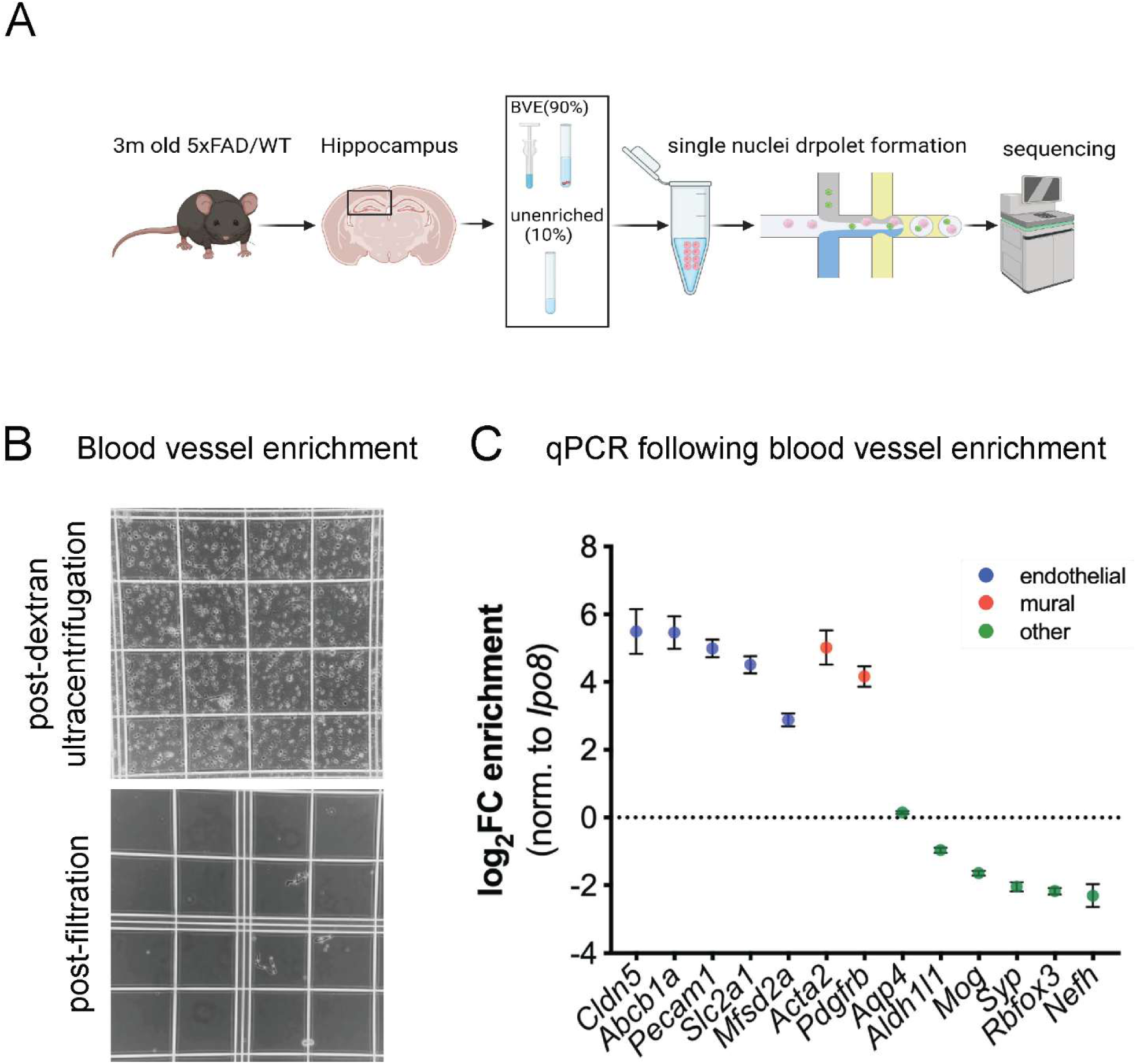
Optimized blood vessel enrichment. (A) Schematic representation of the single­nucleus RNA sequencing (snRNA-Seq) workflow used to analyze the mouse hippocampus. (B) Representative images of mouse cortex homogenates following dextran ultracentrifugation and filtration. Blood vessels are visibly enriched. (C) Quantitative PCR (qPCR) results on n=3 samples of flash-frozen mouse cortex.

**Figure S5.**
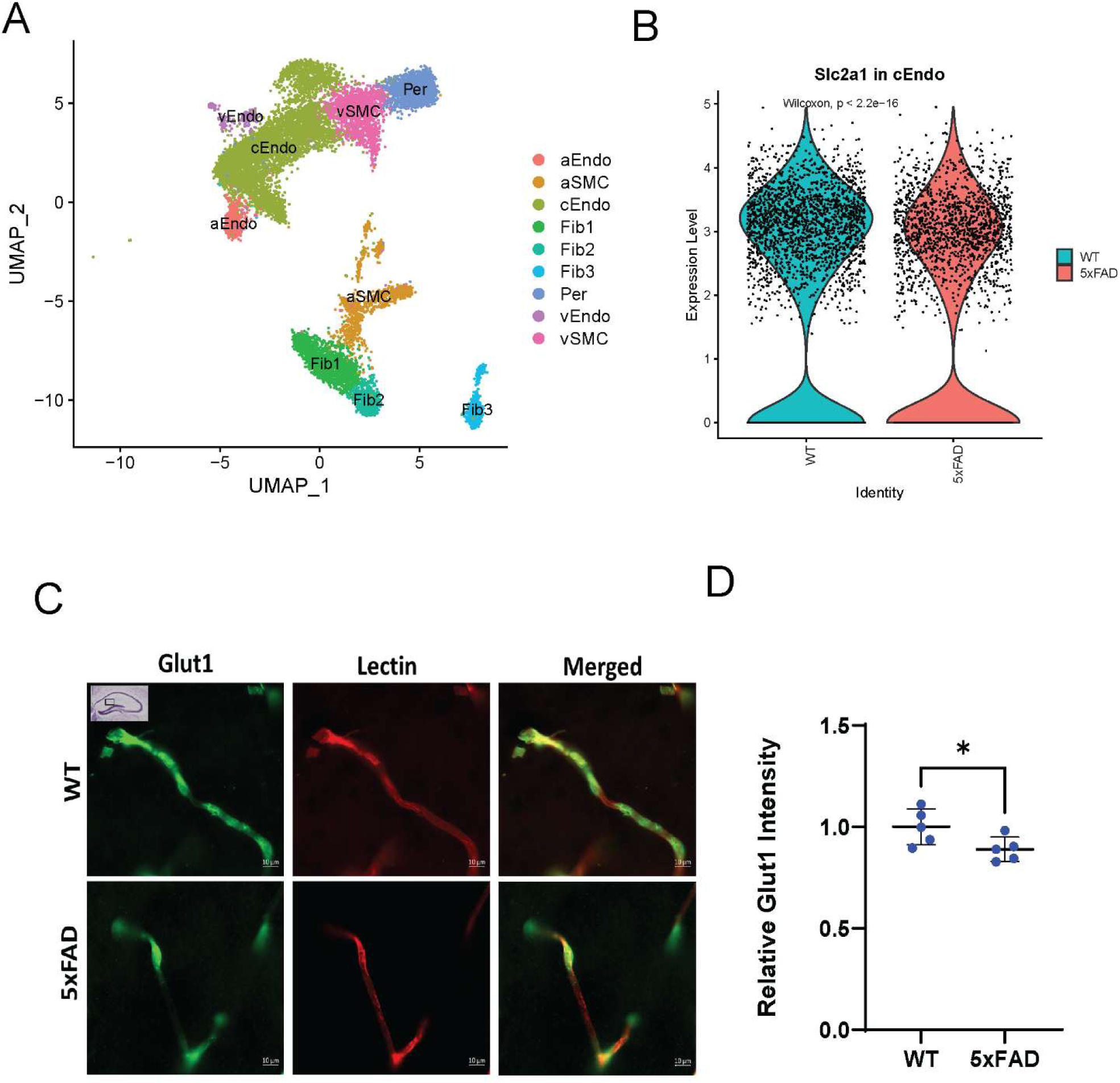
Validation of downregulated Slc2a1 in the blood vessel of prodromal 5XFAD mice. (A) UMAP plot visualizing vascular single nuclei annotated by subtypes. (B) Violin plot shows the significantly reduced mRNA levels of Slc2a1 in the capillary endothelia cells of the hippocampus in 3-month-old 5XFAD mice. (C) Representative immunofluorescent staining of GLUT1, encoded by Slc2a1, demonstrates a decrease in this glucose transporter within cerebral blood vessels of the hippocampus in 3-month-old 5XFAD mice. (D) Quantification of the immunofluorescent intensity of Glut1 (left panel) and Lectin (right panel) in the hippocampal blood vessels. n= 5. *p<0.05 by two-tailed Student’s t-test.

## References

1 Price, J. L. et al. Neuropathology of nondemented aging: presumptive evidence for preclinical Alzheimer disease. Neurobiol Aging 30, 1026–1036, doi:10.1016/j.neurobiolaging.2009.04.002 (2009).

2 Montagne, A. et al. Blood-brain barrier breakdown in the aging human hippocampus. Neuron 85, 296–302, doi: 10.1016/j.neuron.2014.12.032 (2015).

3 Montagne, A., Zhao, Z. & Zlokovic, B. V. Alzheimer’s disease: A matter of blood-brain barrier dysfunction? J Exp Med 214, 3151–3169, doi:10.1084/jem.20171406 (2017).

4 Dickie, B. R., Boutin, H., Parker, G. J. M. & Parkes, L. M. Alzheimer’s disease pathology is associated with earlier alterations to blood-brain barrier water permeability compared with healthy ageing in TgF344-AD rats. NMR Biomed 34, e4510, doi:10.1002/nbm.4510 (2021).

5 Nation, D. A. et al. Blood-brain barrier breakdown is an early biomarker of human cognitive dysfunction. Nat Med 25, 270–276, doi:10.1038/s41591-018-0297-y (2019).

6 van de Haar, H. J. et al. Blood-Brain Barrier Leakage in Patients with Early Alzheimer Disease. Radiology 281, 527–535, doi: 10.1148/radiol.2016152244 (2016).

7 Yang, L., Song, J., Nan, D., Wan, Y. & Guo, H. Cognitive Impairments and blood-brain Barrier Damage in a Mouse Model of Chronic Cerebral Hypoperfusion. Neurochem Res 47, 3817–3828, doi: 10.1007/S11064-022-03799-3 (2022).

8 Geng, J. et al. Blood-Brain Barrier Disruption Induced Cognitive Impairment Is Associated With Increase of Inflammatory Cytokine. Front Aging Neurosci 10, 129, doi: 10.3389/fnagi.2018.00129 (2018).

9 Stranahan, A. M., Hao, S., Dey, A., Yu, X. & Baban, B. Blood-brain barrier breakdown promotes macrophage infiltration and cognitive impairment in leptin receptor-deficient mice. J Cereb Blood Flow Metab 36, 2108–2121, doi: 10.1177/0271678X16642233 (2016).

10 Merlini, M. et al. Fibrinogen Induces Microglia-Mediated Spine Elimination and Cognitive Impairment in an Alzheimer’s Disease Model. Neuron 101, 1099–1108 e1096, doi:10.1016/j.neuron.2019.01.014 (2019).

11 Kim, S., Sharma, C., Jung, U. J. & Kim, S. R. Pathophysiological Role of Microglial Activation Induced by Blood-Borne Proteins in Alzheimer’s Disease. Biomedicines 11, doi: 10.3390/biomedicines11051383 (2023).

12 Ju, F. et al. Increased BBB Permeability Enhances Activation of Microglia and Exacerbates Loss of Dendritic Spines After Transient Global Cerebral Ischemia. Front Cell Neurosci 12, 236, doi: 10.3389/fncel.2018.00236 (2018).

13 Sweeney, M. D., Sagare, A. P. & Zlokovic, B. V. Blood-brain barrier breakdown in Alzheimer disease and other neurodegenerative disorders. Nat Rev Neurol 14, 133–150, doi: 10.1038/nrneurol.2017.188 (2018).

14 Sweeney, M. D., Zhao, Z., Montagne, A., Nelson, A. R. & Zlokovic, B. V. Blood-Brain Barrier: From Physiology to Disease and Back. Physiol Rev 99, 21–78, doi:10.1152/physrev.00050.2017 (2019).

15 Langbaum, J. B. et al. Categorical and correlational analyses of baseline fluorodeoxyglucose positron emission tomography images from the Alzheimer’s Disease Neuroimaging Initiative (ADNI). Neuroimage 45, 1107–1116, doi:10.1016/j.neuroimage.2008.12.072 (2009).

16 Hoyer, S., Nitsch, R. & Oesterreich, K. Predominant abnormality in cerebral glucose utilization in late-onset dementia of the Alzheimer type: a cross-sectional comparison against advanced late-onset and incipient early-onset cases. J Neural Transm Park Dis Dement Sect 3, 1–14, doi:10.1007/BF02251132 (1991).

17 Fukuyama, H. et al. Altered cerebral energy metabolism in Alzheimer’s disease: a PET study. J Nucl Med 35, 1–6(1994).

18 Vaishnavi, S. N. et al. Regional aerobic glycolysis in the human brain. Proc Natl Acad Sci USA 107, 17757–17762, doi: 10.1073/pnas.1010459107 (2010).

19 Vlassenko, A. G. et al. Spatial correlation between brain aerobic glycolysis and amyloid-beta (Abeta) deposition. Proc Natl Acad Sci U S A 107, 17763–17767, doi:10.1073/pnas.1010461107 (2010).

20 Hu, Y. et al. mTOR-mediated metabolic reprogramming shapes distinct microglia functions in response to lipopolysaccharide and ATP. Glia 68, 1031–1045, doi:10.1002/glia.23760 (2020).

21 Oakley, H. et al. Intraneuronal beta-amyloid aggregates, neurodegeneration, and neuron loss in transgenic mice with five familial Alzheimer’s disease mutations: potential factors in amyloid plaque formation. J Neurosci 26, 10129–10140, doi:10.1523/JNEUROSCI.1202-06.2006 (2006).

22 Padua, M. S., Guil-Guerrero, J. L. & Lopes, P. A. Behaviour Hallmarks in Alzheimer’s Disease 5xFAD Mouse Model. Int J Mol Sci 25, doi:10.3390/ijms25126766 (2024).

23 Takechi, R. et al. Blood-Brain Barrier Dysfunction Precedes Cognitive Decline and Neurodegeneration in Diabetic Insulin Resistant Mouse Model: An Implication for Causal Link. Front Aging Neurosci 9, 399, doi:10.3389/fnagi.2017.00399 (2017).

24 Wei, Z. et al. Non-contrast assessment of blood-brain barrier permeability to water in mice: An arterial spin labeling study at cerebral veins. Neuroimage 268, 119870, doi: 10.1016/j.neuroimage.2023.119870 (2023).

25 Lin, Z. et al. Non-contrast MR imaging of blood-brain barrier permeability to water. Magn Reson Med 80, 1507–1520, doi:10.1002/mrm.27141 (2018).

26 Lin, Z. et al. Blood-Brain Barrier Breakdown in Relationship to Alzheimer and Vascular Disease. Ann Neurol 90, 227–238, doi:10.1002/ana.26134 (2021).

27 Zhukov, O. et al. Preserved blood-brain barrier and neurovascular coupling in female 5xFAD model of Alzheimer’s disease. Front Aging Neurosci 15, 1089005, doi: 10.3389/fnagi.2023.1089005 (2023).

28 Hock, C. et al. Decrease in parietal cerebral hemoglobin oxygenation during performance of a verbal fluency task in patients with Alzheimer’s disease monitored by means of near-infrared spectroscopy (NIRS)­correlation with simultaneous rCBF-PET measurements. Brain Res 755, 293–303, doi: 10.1016/s0006-8993(97)00122-4 (1997).

29 Iturria-Medina, Y. et al. Early role of vascular dysregulation on late-onset Alzheimer’s disease based on multifactorial data-driven analysis. Nat Commun 7, 11934, doi: 10.1038/ncommsl 1934 (2016).

30 Wise, R. G., Harris, A. D., Stone, A. J. & Murphy, K. Measurement of OEF and absolute CMRO2: MRI-based methods using interleaved and combined hypercapnia and hyperoxia. Neuroimage 83, 135–147, doi: 10.1016/j.neuroimage.2013.06.008 (2013).

31 Wei, Z. et al. Quantitative assessment of cerebral venous blood T(2) in mouse at 11.7T: Implementation, optimization, and age effect. Magn Reson Med 80, 521–528, doi:10.1002/mrm.27046 (2018).

32 ladecola, C. Neurovascular regulation in the normal brain and in Alzheimer’s disease. Nat Rev Neurosci 5, 347–360, doi:10.1038/nrn1387 (2004).

33 Sun, X. et al. Hypoxia facilitates Alzheimer’s disease pathogenesis by up-regulating BACE1 gene expression. Proc Natl Acad Sci U S A 103, 18727–18732, doi:10.1073/pnas.0606298103 (2006).

34 Preece, P. & Cairns, N. J. Quantifying mRNA in postmortem human brain: influence of gender, age at death, postmortem interval, brain pH, agonal state and inter-lobe mRNA variance. Brain Res Mol Brain Res 118, 60–71, doi:10.1016/s0169-328x(03)00337-1 (2003).

35 Hagihara, H. et al. Large-scale animal model study uncovers altered brain pH and lactate levels as a transdiagnostic endophenotype of neuropsychiatric disorders involving cognitive impairment. Elife 12, doi: 10.7554/eLife.89376 (2024).

36 Decker, Y. et al. Decreased pH in the aging brain and Alzheimer’s disease. Neurobiol Aging 101, 40–49, doi: 10.1016/j.neurobiolaging.2020.12.007 (2021).

37 Schwartz, L., Peres, S., Jolicoeur, M. & da Veiga Moreira, J. Cancer and Alzheimer’s disease: intracellular pH scales the metabolic disorders. Biogerontology 21, 683–694, doi: 10.1007/s10522-020-09888-6 (2020).

38 Lyros, E. et al. Normal brain aging and Alzheimer’s disease are associated with lower cerebral pH: an in vivo histidine (1)H-MR spectroscopy study. Neurobiol Aging 87, 60–69, doi: 10.1016/j.neurobiolaging.2019.11.012 (2020).

39 Gonzales, E. B. & Sumien, N. Acidity and Acid-Sensing Ion Channels in the Normal and Alzheimer’s Disease Brain. J Alzheimers Dis 57, 1137–1144, doi:10.3233/JAD-161131 (2017).

40 Chen, L. et al. Early detection of Alzheimer’s disease using creatine chemical exchange saturation transfer magnetic resonance imaging. Neuroimage 236, 118071, doi: 10.1016/j.neuroimage.2021.118071 (2021).

41 Zhang, D. et al. Metabolic regulation of gene expression by histone lactylation. Nature 574, 575­-580, doi: 10.1038/S41586-019-1678-1 (2019).

42 Pan, R. Y. et al. Positive feedback regulation of microglial glucose metabolism by histone H4 lysine 12 lactylation in Alzheimer’s disease. Cell Metab 34, 634–648 e636, doi: 10.1016/j.cmet.2022.02.013 (2022).

43 Garcia, F. J. et al. Single-cell dissection of the human brain vasculature. Nature 603, 893–899, doi: 10.1038/S41586-022-04521-7 (2022).

44 Sun, N. et al. Single-nucleus multiregion transcriptomic analysis of brain vasculature in Alzheimer’s disease. Nat Neurosci 26, 970–982, doi: 10.1038/s41593-023-01334-3 (2023).

45 Tang, X. et al. Spatial learning and memory impairments are associated with increased neuronal activity in 5XFAD mouse as measured by manganese-enhanced magnetic resonance imaging. Oncotarge*t7*, 57556–57570, doi:10.18632/oncotarget.11353 (2016).

46 Gu, L. et al. Myelin changes at the early stage of 5XFAD mice. Brain Res Bull 137, 285–293, doi:10.1016/j.brainresbull.2017.12.013 (2018).

47 Selak, S. et al. A role for SNAP25 in internalization of kainate receptors and synaptic plasticity. Neuron 63, 357–371, doi:10.1016/j.neuron.2009.07.017 (2009).

48 Zhang, C., Xie, S. & Malek, M. SNAP-25: A biomarker of synaptic loss in neurodegeneration. Clin ChimActa 571, 120236, doi:10.1016/j.cca.2025.120236 (2025).

49 Coba, M. P. et al. Dlgapl knockout mice exhibit alterations of the postsynaptic density and selective reductions in sociability. Sci Rep 8, 2281, doi:10.1038/s41598-018-20610-y (2018).

50 Rasmussen, A. H., Rasmussen, H. B. & Silahtaroglu, A. The DLGAP family: neuronal expression, function and role in brain disorders. Mol Brain 10, 43, doi: 10.1186/s13041-017-0324-9 (2017).

51 Giannoni, P. et al. Cerebrovascular pathology during the progression of experimental Alzheimer’s disease. Neurobiol Dis 88, 107–117, doi: 10.1016/j.nbd.2O16.01.001 (2016).

52 Correia, S. C. & Moreira, P. I. Oxygen Sensing and Signaling in Alzheimer’s Disease: A Breathtaking Story! Cell Mol Neurobiol 42, 3–21, doi: 10.1007/s10571-021-01148-6 (2022).

53 Di Marco, L. Y. et al. Vascular dysfunction in the pathogenesis of Alzheimer’s disease--A review of endothelium-mediated mechanisms and ensuing vicious circles. Neurobiol Dis 82, 593–606, doi:10.1016/j.nbd.2015.08.014 (2015).

54 Ostergaard, L. et al. The capillary dysfunction hypothesis of Alzheimer’s disease. Neurobiol Aging 34, 1018–1031, doi: 10.1016/j.neurobiolaging.2012.09.011 (2013).

55 Wang, W. et al. Identification of early Alzheimer’s disease subclass and signature genes based on PANoptosis genes. Front Immunol 15, 1462003, doi: 10.3389/fimmu.2024.1462003 (2024).

56 Wesenhagen, K. E. J. et al. Synaptic protein CSF levels relate to memory scores in individuals without dementia. Alzheimers Res Therî7, 56, doi: 10.1186/s13195-025-01703-z (2025).

57 Duits, F. H. et al. Serial Cerebrospinal Fluid Sampling Reveals Trajectories of Potential Synaptic Biomarkers in Early Stages of Alzheimer’s Disease. J Alzheimers Dis 100, S103–S114, doi:10.3233/JAD-240610 (2024).

58 Wei, Z. et al. Brain metabolism in tau and amyloid mouse models of Alzheimer’s disease: An MRI study. NMR Biomed 34, e4568, doi: 10.1002/nbm.4568 (2021).

59 Wei, Z. et al. Optimization of phase-contrast MRI for the estimation of global cerebral blood flow of mice at 11,7T. Magn Reson Med 81, 2566–2575, doi: 10.1002/mrm.27592 (2019).

60 Chen, L., et al. High-resolution creatine mapping of mouse brain at 11.7 T using non-steady-state chemical exchange saturation transfer. NMR Biomed 32, e4168, doi:10.1002/nbm.4168 (2019).

61 Chen, L., et al. Early detection of Alzheimer’s disease using creatine chemical exchange saturation transfer magnetic resonance imaging. Neuroimage 236, 118071 (2021).

62 Lee, Y. K., Uchida, H., Smith, H., Ito, A. & Sanchez, T. The isolation and molecular characterization of cerebral microvessels. Nat Protoc 14, 3059–3081, doi:10.1038/s41596-019-0212-0 (2019).

63 Lee, H. et al. Cell Type-Specific Transcriptomics Reveals that Mutant Huntingtin Leads to Mitochondrial RNA Release and Neuronal Innate Immune Activation. Neuron 107, 891–908 e898, doi: 10.1016/j.neuron.2020.06.021 (2020)

